# Tomosyn-2 Regulates Postnatal β-Cell Expansion and Insulin Secretion to Maintain Glucose Homeostasis

**DOI:** 10.1101/2025.05.19.654959

**Authors:** Katherine C. Perez, Justin Alexander, Md Mostafizur Rahman, Haifa A. Alsharif, Yanping Liu, Jeong-A Kim, Chad S. Hunter, Thanh Nguyen, Sushant Bhatnagar

## Abstract

The transition from proliferative to functionally mature β-cells is a critical developmental process, yet the molecular mechanisms that coordinate this shift remain poorly understood. Here, we identify Tomosyn-2 as a key regulator of β-cell maturation. Tomosyn-2 expression declines with age in mouse islets, coinciding with enhanced biphasic glucose-stimulated insulin secretion (GSIS) and reduced β-cell proliferation. Genetic deletion of Tomosyn-2 improves glucose tolerance, elevates plasma insulin levels, and augments islet insulin secretion, without altering systemic insulin sensitivity. Mechanistically, Tomosyn-2 interacts with syntaxin-1A (Stx1A) to inhibit insulin granule exocytosis by limiting SNARE complex formation. Transcriptomic and network analyses reveal that Tomosyn-2 loss reprograms gene expression to strengthen the coupling between insulin secretion and proliferative pathways. Its deletion also reduces β-cell proliferation and mass expansion, suppresses cell cycle and Akt1 signaling, and promotes β-cell identity, maturation, and altered islet architecture. These findings identify Tomosyn-2 as a crucial molecular switch that orchestrates the balance between proliferation and functional maturation during postnatal β-cell development.

## Introduction

The transition from immature to functionally mature pancreatic β-cells is a tightly regulated developmental process defined by increased glycolysis, mitochondrial oxidative phosphorylation, insulin granule trafficking, insulin exocytotic machinery, and glucose-stimulated insulin secretion (GSIS)^1–4^. Postnatally, immature β-cells continue to proliferate to establish a critical mass of mature β-cells required for sustained insulin release and the maintenance of glucose homeostasis^1,5–7^. Proliferating, immature β-cells are characterized by the expression of immaturity markers such as hexokinases (Hk 1–3), elevated basal insulin secretion, and blunted GSIS^8,9^. During the transition from neonatal to adult stages, the proportion of proliferating β-cells declines as functionally mature β-cells accumulate^10^, highlighting an inverse relationship between proliferative capacity and functional maturity. Proper coordination of these processes during postnatal development is essential, as disruption can lead to long-term metabolic dysfunction^11,12^. Inadequate β-cell mass expansion or aberrant insulin secretion increases susceptibility to β-cell failure and the development of type 2 diabetes (T2D) in the context of aging or obesity^13–16^. Elucidating the molecular mechanisms that govern the dynamic balance between β-cell proliferation and functional maturation is therefore critical for developing therapeutic strategies to preserve β-cell mass and function in metabolic disease. β-cell functional maturation is orchestrated by a coordinated transcriptional program that is dynamically affected by nutritional cues^1,17–19^. This maturation trajectory, culminating in regulated insulin secretion, is modulated by a network of β-cell lineage-defining transcription factors (e.g., *Pdx1, NeuroD1, Nkx6.1*), nutrient-responsive regulators (e.g., *MafA, Foxo1, NFATc1/2, CREB*), and metabolic signaling genes (e.g., *Glut2, Gck, Glp1R*)^20^. Nkx6.1 promotes the expression of *Glut2* and *Glp1R*, while Pdx1 regulates MafA and genes essential for insulin biosynthesis (*Ins1*, *Ins2*, *Pcsk1*, *Pcsk2*) and secretion (e.g., syntaxins, synaptotagmins)^9,21^. MafA, in turn, cooperates with Pdx1 and NeuroD1 to activate *Urocortin-3* (*Ucn3*), a key marker of β-cell maturity whose expression peaks during early postnatal development and correlates with β-cell functional competence^22^. Loss of Ucn3 leads to β-cell dedifferentiation, marked by the re-expression of “disallowed” genes such as Hk1 and Ldha^1,23^. Despite these insights, how nutritional signals—particularly glucose—interface with transcriptional networks during the critical post-weaning period remains poorly understood.

Lineage-tracing studies in mice have shown that replicating pre-existing β-cells primarily drive postnatal expansion of β-cell mass^24–26^. Several key pathways regulate this proliferation response, including insulin, IGF1, mTOR, and Wnt/β-catenin signaling^6^. Among these, the IRS2–PI3K–Akt1 axis plays a central role in promoting β-cell survival and proliferation by modulating key effectors, such as Foxo1, Gsk3β, p27Kip1 (Cdkn1b), and Cyclin D1^27–31^. Genetic ablation of IRS2 leads to impaired β-cell proliferation and reduced β-cell mass, underscoring its critical role^32,33^. However, chronic insulin hypersecretion may attenuate IRS2–Akt1 signaling via negative feedback^27,34^, potentially suppressing proliferation. The direct contribution of β-cell-derived insulin to β-cell proliferation remains unclear^35,36^. Moreover, sustained insulin production can induce ER stress and activate the unfolded protein response (UPR)—including sXbp1, Atf4, and Bip—further impairing β-cell proliferation and contributing to neonatal diabetes^5,37^. Thus, the molecular mechanisms linking insulin secretion with β-cell proliferation remain insufficiently elucidated.

A defining feature of mature β-cells is their ability to execute biphasic insulin secretion in response to glucose—a rapid first phase followed by a sustained second phase—critical for glucose homeostasis^38,39^. Insulin is stored in granules that undergo trafficking, docking, and fusion with the plasma membrane (PM) in a process mediated by the SNARE (soluble N-ethylmaleimide-sensitive factor attachment protein receptor) complex^40–43^. Core SNARE components—syntaxins (Stx1A–4), VAMP2/8, and SNAP25/23—assemble into a four-helix bundle to drive membrane fusion^44–46^. The efficiency of SNARE-mediated fusion is fine-tuned by accessory proteins that either facilitate or inhibit complex formation, thereby imparting precise control of insulin secretion^46–51^. Thus, SNARE complex formation is a critical checkpoint for insulin release kinetics^52^. Tomosyn-2, initially characterized in neuroendocrine cells, has emerged as a conserved inhibitor of vesicle exocytosis^53–55^. It contains an N-terminal WD40 repeat domain and a C-terminal VAMP-like motif but lacks a transmembrane domain^56,57^. Functionally, Tomosyn-2 acts as an endogenous inhibitor of SNARE complex formation and insulin secretion^48,58^.

Tomosyn-2 was positionally cloned under a fasting glucose quantitative trait locus (QTL) in an F2 intercross between nondiabetic-obese and diabetic-obese mouse strains. A non-synonymous coding variant (S912L) in Tomosyn-2 enhanced its protein stability and was associated with impaired insulin secretion, hypoinsulinemia, and hyperglycemia^59^. Tomosyn-2 levels are regulated post-translationally; glucose and other insulin secretagogues promote its phosphorylation and degradation, thereby relieving Tomosyn-2’s inhibitory effect on insulin secretion^48^. Nevertheless, the physiological role of Tomosyn-2 in β-cell development, identity, and systemic glucose homeostasis remains largely unknown.

Given the critical need to balance β-cell proliferation and maturation for optimal β-cell mass and metabolic function, we investigated the role of Tomosyn-2 in postnatal β-cell development. Here, we identify Tomosyn-2 as a key regulator that reciprocally controls β-cell proliferation and functional maturation. Loss of Tomosyn-2 enhances β-cell identity, insulin secretory capacity, and maturation markers while suppressing proliferation, ultimately shaping islet architecture, β-cell mass, and whole-body glucose homeostasis during postnatal development.

## Results

### Progressive enhancement of biphasic insulin secretion inversely correlates with β-cell proliferation during neonatal to young adult islet development

Postnatal β-cell proliferation is crucial for establishing sufficient β-cell mass to meet metabolic demands in adulthood^1^. To assess the relationship between insulin secretion and β-cell proliferation during postnatal development, we measured biphasic GSIS from islets of C57BL/6J mice at 1, 2, 4, 6, and 14 weeks of age.

Insulin secretion was determined in response to low (2.8 mM) and high (16.7 mM) glucose concentrations (Fig. 1A–F, Supplementary Fig. 1). Basal insulin secretion at 2.8 mM glucose remained unchanged across 2-to 14-week-old age groups (Fig. 1A), whereas high glucose stimulation elicited a biphasic insulin secretion response in all groups (Fig. 1A) with significantly enhanced responses in islets from 4-, 6-, and 14-week-old mice compared to 1- and 2-week-old neonates (Fig. 1B–C). Quantification of the area under the curve (AUC) for the first (6–11 min) and sustained (12–45 min) phases of insulin secretion, both of which progressively increased with age. Although secretion kinetics were comparable between 1- and 2-week-old islets, distinct stepwise increases were observed at key developmental transitions: 4 vs. 2 weeks, 6 vs. 4 weeks, and 14 vs. 6 weeks (Fig. 1E–F, Supplementary Fig. 1H–1I), showing a linear increase in the early and sustained phases of GSIS from neonatal to young adult stages. These trends were consistent whether insulin output was normalized to total insulin content (Fig. 1A–C), basal insulin secretion at 2.8 mM glucose (Supplementary Fig. 1H–1I), or islet number (Supplementary Fig. 1A–G), implicating a progressive functional maturation of β-cells from neonatal to young adult stages.

**Figure 1.**
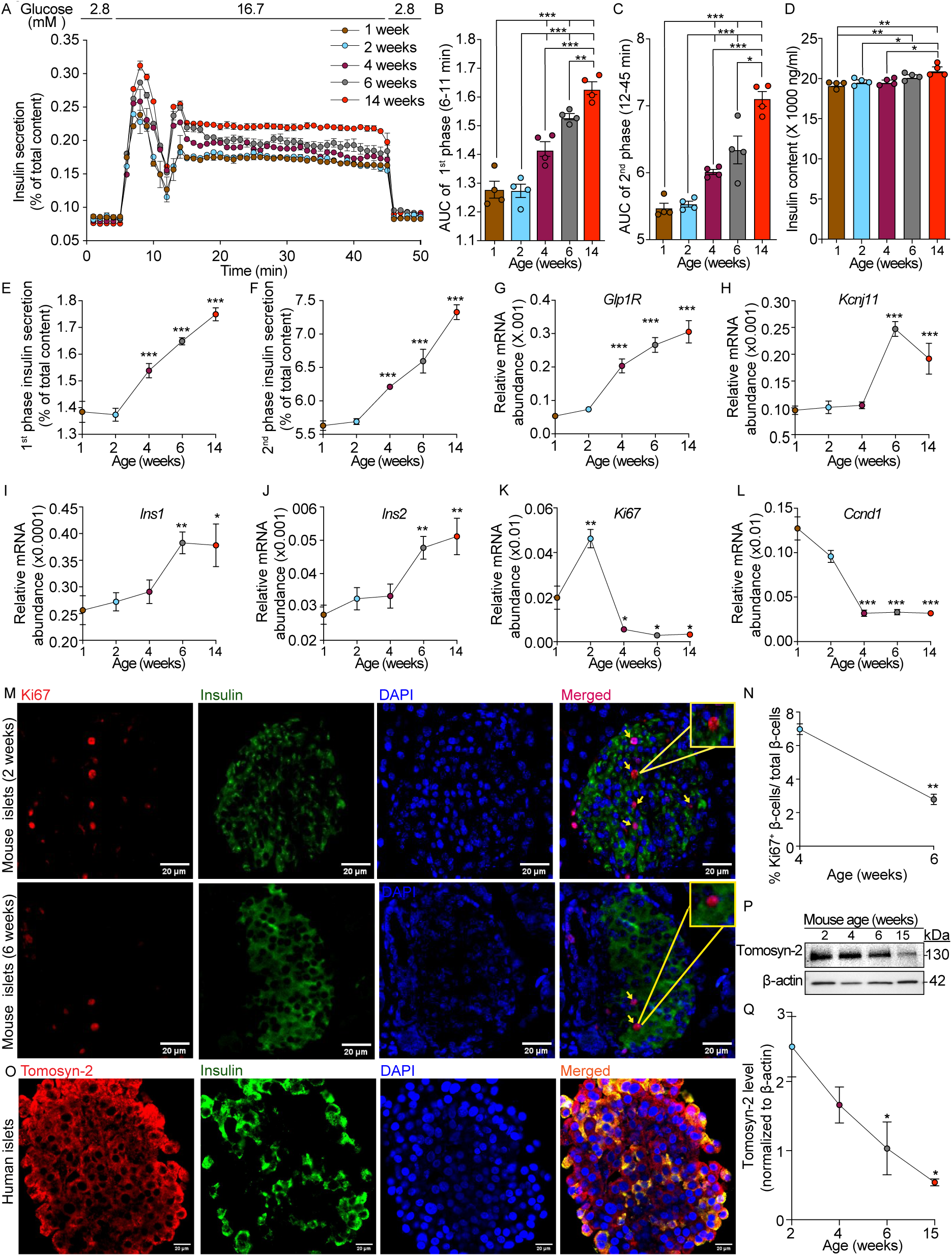
Age-dependent regulation of insulin secretion, β-cell proliferation, and Tomosyn-2 expression in C57BL/6J (B6) mouse islets. (A) Dynamic glucose-stimulated insulin secretion (GSIS) from isolated islets of 1-, 2-, 4-, 6-, and 14-week-old B6 mice exposed sequentially to 2.8 mM and 16.7 mM glucose. *N = 4 biological replicates per group.* Quantification of the area under the curve (AUC) for the (B) first phase of insulin secretion (6–11 min) and (C) AUC second phase (12–45 min). (D) Total insulin content of islets from each age group. (E, F) GSIS during the first (E) and second (F) phases in response to 16.7 mM glucose. (G–L) mRNA expression levels of key β-cell genes in islets from B6 mice at 1, 2, 4, 6, and 14 weeks of age:(G) Glp1r, (H) Kcnj11, (I) Ins1, (J) Ins2, (K) Ki67, and (L) Ccnd1. (M) IF staining of insulin (green) and Ki67 (red) in islets from 4-week-old (top panel) and 6-week-old (bottom panel) B6 mice. Arrows indicate Ki67⁺β-cells; insets show magnified views of highlighted regions. (N) Quantification of Ki67⁺ β-cell percentage in islets from 2-versus 6-week-old mice (n = 3; ∼200 islets counted per group). (O) IF staining for Tomosyn-2 (red) and insulin (green) in human islets (donors ND1547, ND1770, ND8316), imaged at 40X magnification. (P) Representative Western blot showing Tomosyn-2 and β-actin protein levels in islets from 1-, 2-, 4-, 6-, and 14-week-old B6 mice (n = 4). (Q) Densitometric quantification of Tomosyn-2 protein expression from (P). Data are presented as mean ± s.e.m. An unpaired two-tailed Student’s t-test determined statistical significance. ^∗^P < 0.05, ^∗∗^P < 0.01, ^∗∗∗^P < 0.001

Corroborating with the increased insulin secretion, insulin protein levels (Fig. 1D) and the mRNA abundance of key genes involved insulin secretion, including *Glp1R*, *Kcnj11, Ins1,* and *Ins2* (Fig. 1G–J), were significantly upregulated in islets from 6- and 14-week-old mice compared to neonates (Fig. 1G–J). Interestingly, when normalized to islet number, total islet insulin content was slightly lower in young adult islets than in neonates (Supplementary Fig. 1D–G), suggesting a functional rather than content-based effect. These data show progressive increases in insulin secretion response to glucose stimulation from neonatal to young adult mouse islets, supporting the functional maturation of β-cells.

To assess whether this functional maturation was associated with changes in β-cell proliferation, we analyzed proliferation markers across the same developmental stages. Gene expression analysis showed a significant decline in proliferation markers such as *Ki67* and *Ccnd1* in islets from 4-, 6-, and 14-week-old mice compared to neonates (Fig. 1K–L). Expression of the apoptosis-related gene *Bcl2* also declined with age (Supplementary Fig. 2A), with no evidence of compensatory cell death. Expression of mitochondrial genes remained unchanged (Supplementary Fig. 2B), ruling out metabolic dysfunction. Immunofluorescence (IF) staining of pancreatic sections revealed a marked age-dependent decline in Ki67^+^β-cells (Fig. 1M–N). Quantitative analysis showed that approximately 7% of β-cells were proliferating at 4 weeks, whereas this proportion dropped to less than 4% by 6 weeks of age (Fig. 1N), consistent with previous reports^25,60^. A similar decline was observed in proliferating non-β-cells (Supplementary Fig. 2).

Together, these findings reveal an inverse relationship between β-cell proliferation and glucose-stimulated insulin secretion at both functional and gene expression levels, implicating a developmental switch in which proliferative β-cells transition into a functionally mature state, which is crucial for establishing an optimal β-cell mass and functional capacity to support glucose homeostasis in young adulthood.

### Age-dependent decline of Tomosyn-2 levels in neonatal to young adult islets

Tomosyn-2, an endogenous inhibitor of insulin secretion^48,61^, is expressed in pancreatic β-cells. IF staining of human islets revealed strong Tomosyn-2 expression (red) in insulin^+^ β-cells (green) (Fig. 1O). In addition to β-cells, Tomosyn-2 was also detectable in glucagon^+^α-cells and somatostatin^+^δ-cells (Supplementary Fig. 4). We previously demonstrated that high glucose stimulation promotes Tomosyn-2 degradation in β-cells^48^, and that a non-synonymous coding variant (S912L) enhances its protein stability, impairing insulin secretion^59^. Building on these findings and considering the progressive increase in GSIS during postnatal development, we hypothesized that Tomosyn-2 expression is dynamically regulated by nutritional cues during this critical period of maturation.

To test this, we quantified Tomosyn-2 protein levels in islets isolated from mice at postnatal weeks 1, 2, 4, 6, and 14. Western blot analysis revealed a progressive, age-dependent decline in Tomosyn-2 expression (Fig. 1P–Q). Compared to week 1, Tomosyn-2 levels decreased by approximately 60% at week 6 and nearly 80% by week 14. This decline occurred in parallel with the previously observed reduction in β-cell proliferation and the enhancement of biphasic GSIS. These findings indicate that Tomosyn-2 is developmentally downregulated during postnatal β-cell maturation. The temporal decline in Tomosyn-2 may function as a regulatory switch, relieving its inhibitory effects on insulin secretion while attenuating proliferative signals—thus facilitating the transition from a proliferative, immature β-cell state to a functionally mature phenotype capable of sustaining glucose homeostasis in adulthood.

### The loss of Tomosyn-2 improves glucose clearance without affecting insulin action in young mice

To investigate the functional and metabolic role of Tomosyn-2 in vivo, we generated Tomosyn-2 deletion (*Tomosyn-2^-/-^)* mice on a C57BL/6J background. IF staining and Western blot analysis confirmed complete loss of Tomosyn-2 expression in islet β-cells (Fig. 2A–C), validating the knockout model for further physiological assessment.

**Figure 2.**
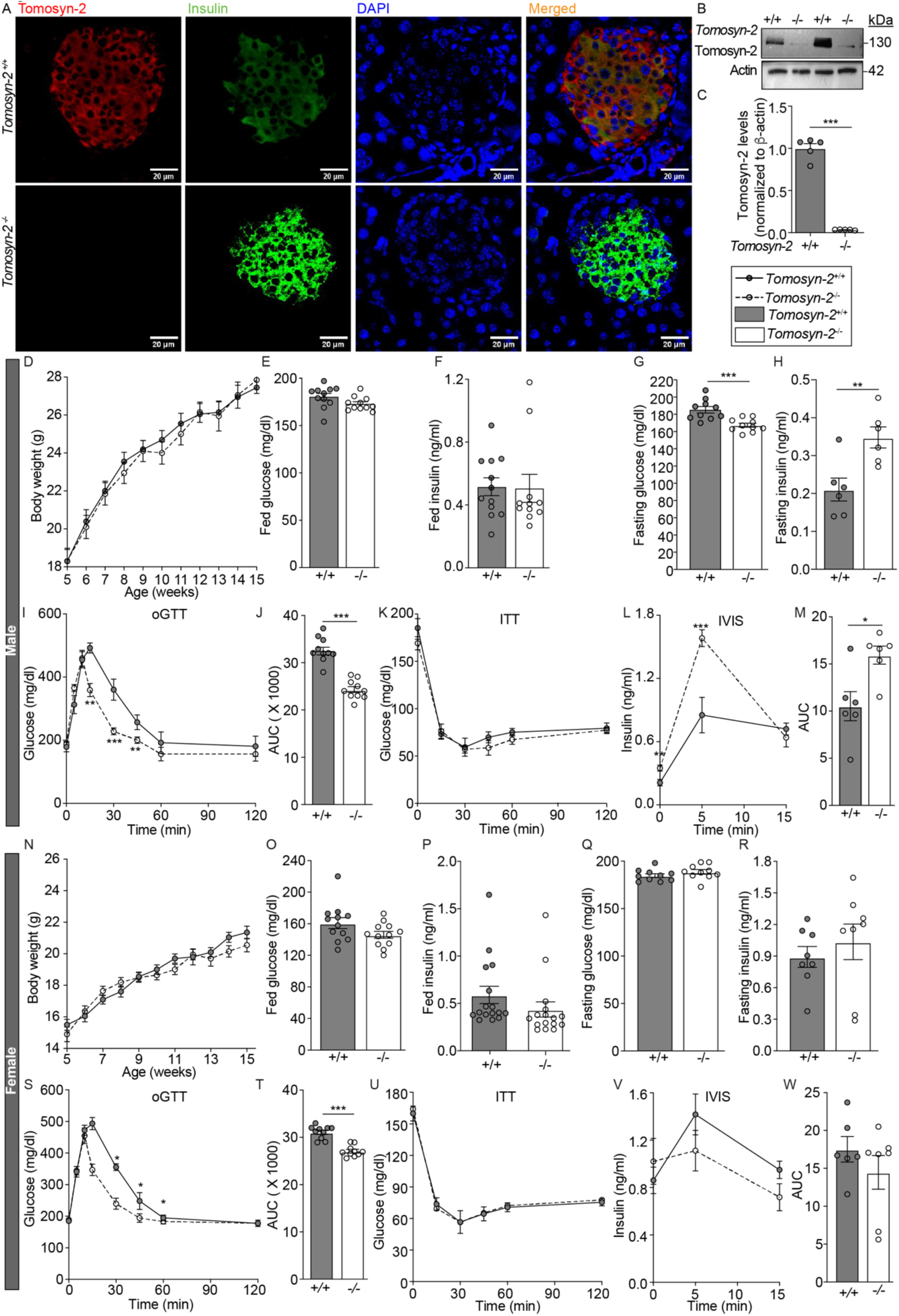
Loss of Tomosyn-2 improves glucose clearance in mice. (A) Representative IF image showing Tomosyn-2 (red) and insulin (green) staining in pancreatic islets from *Tomosyn-2^-/-^* and *Tomosyn-2^+/+^* mice (20X magnification, n = 4). (B) Western blot analysis of Tomosyn-2 and β-actin protein levels in isolated islets from *Tomosyn-2^-/-^*and *Tomosyn-2^+/+^* mice (n > 5). (C) Relative *Tomosyn-2* mRNA expression in islets of *Tomosyn-2^-/-^* and *Tomosyn-2^+/+^* mice (n = 5). (D) Body weight of male *Tomosyn-2^-/-^* and *Tomosyn-2^+/+^* mice from 5-15 weeks of age (n > 20). (E–F) Fed blood glucose (E) and plasma insulin (F) levels measured at 8 A.M. in 6-week-old male mice (n > 10). (G–H) Fasting (6 h) blood glucose (G) and plasma insulin (H) levels in 6-week-old male mice (n > 10). (I) Oral glucose tolerance test (OGTT) was performed in 6-week-old male mice after 6 h of fasting (n = 10). (J) Quantification of glucose area under the curve (AUC) during OGTT from (I). (K) Insulin tolerance test (ITT) was performed by intraperitoneal injection of 0.5 U/kg human insulin in 6-week-old male mice after 6 h fasting (n > 6). (L–M) Plasma insulin levels (L) and corresponding AUC (M) in response to OGTT (n = 6). (N) Body weight of female *Tomosyn-2^-/-^* and *Tomosyn-2^+/+^* mice from 5 to 15 weeks of age (n > 20). (O–P) Fed blood glucose (O) and plasma insulin (P) levels in 6-week-old female mice (n > 10). (Q–R) Fasting (6 h) blood glucose (Q) and plasma insulin (R) levels in 6-week-old female mice (n = 8). (S) OGTT in 6-week-old female mice after 6 h fasting (n = 10). (T) The glucose AUC from the OGTT is shown in (S). (U) ITT was performed in 6-week-old female mice as described in (K) (n > 6). (V–W) Plasma insulin levels (V) and AUC (W) during OGTT in female mice (n = 6). Data are presented as mean ± s.e.m. An unpaired two-tailed Student’s t-test determined statistical significance. ^∗^P < 0.05, ^∗∗^P < 0.01, ^∗∗∗^P < 0.001.

We first examined whole-body glucose homeostasis in *Tomosyn-2^-/-^*mice. Age-dependent increases in body weight (BW) were observed in control male and female mice, while Tomosyn-2 deletion did not affect BW in either sex (Fig. 2D, 2N). In male *Tomosyn-2^-/-^* mice, fasting (6 h) plasma glucose levels were significantly reduced (p < 0.001), accompanied by elevated fasting plasma insulin levels (p < 0.01), compared to *Tomosyn-2^+/+^* littermates (Fig. 2G–H). In contrast, fed glucose and insulin levels were unchanged between genotypes (Fig. 2E–F), indicating that Tomosyn-2 loss predominantly affects fasting metabolic parameters.

We performed oral glucose tolerance tests (OGTT) in male *Tomosyn-2^-/-^* and *Tomosyn- 2^+/+^* mice to examine glucose clearance. Tomosyn-2 deletion resulted in significantly improved glucose clearance, with a ∼25% reduction in glucose AUC (AUC; p < 0.0001) compared to controls (Fig. 2I–J). However, insulin tolerance tests (ITT) revealed no significant differences in insulin action between groups (Fig. 2K), suggesting that the improved glucose tolerance was not due to insulin action.

To directly assess insulin secretory function in vivo, we measured plasma insulin levels following oral glucose administration after a 6-hour fast. Both genotypes exhibited a glucose-induced rise in plasma insulin at 5 and 15 minutes; however, *Tomosyn-2^-/-^* male mice showed a significantly greater increase (∼50%, p = 0.014) in plasma insulin levels as measured by AUC, compared to *Tomosyn-2^+/+^* controls (Fig. 2L–M). These findings indicate that the improved glucose clearance in *Tomosyn-2^-/-^* mice is primarily driven by enhanced insulin secretion rather than altered peripheral insulin action.

A similar trend was observed in female *Tomosyn-2^-/-^* mice, which also exhibited improved glucose clearance without changes in insulin sensitivity. However, unlike males, female deletion mice did not display elevated fasting insulin levels or increased glucose-stimulated insulin secretion, suggesting a sex-dependent divergence in the physiological response to Tomosyn-2 loss. Together, these results show that Tomosyn-2 negatively regulates insulin secretion in vivo and that its genetic ablation enhances glucose-stimulated insulin release, thereby improving glucose homeostasis, particularly in male mice.

### Loss of Tomosyn-2 increases biphasic glucose-stimulated insulin secretion

To determine the role of Tomosyn-2 in stimulus-coupled insulin secretion, we performed perifusion assays using size-matched islets (n = 80) isolated from 6-week-old male and female *Tomosyn-2^-/-^* and *Tomosyn-2^+/+^* mice. Islets were sequentially perifused with low glucose (2.8 mM), high glucose (16.7 mM), returned to basal glucose, and finally depolarized with potassium chloride (KCl; 40 mM), to assess both glucose- and depolarization-induced insulin release (Fig. 3A, 3J).

**Figure 3.**
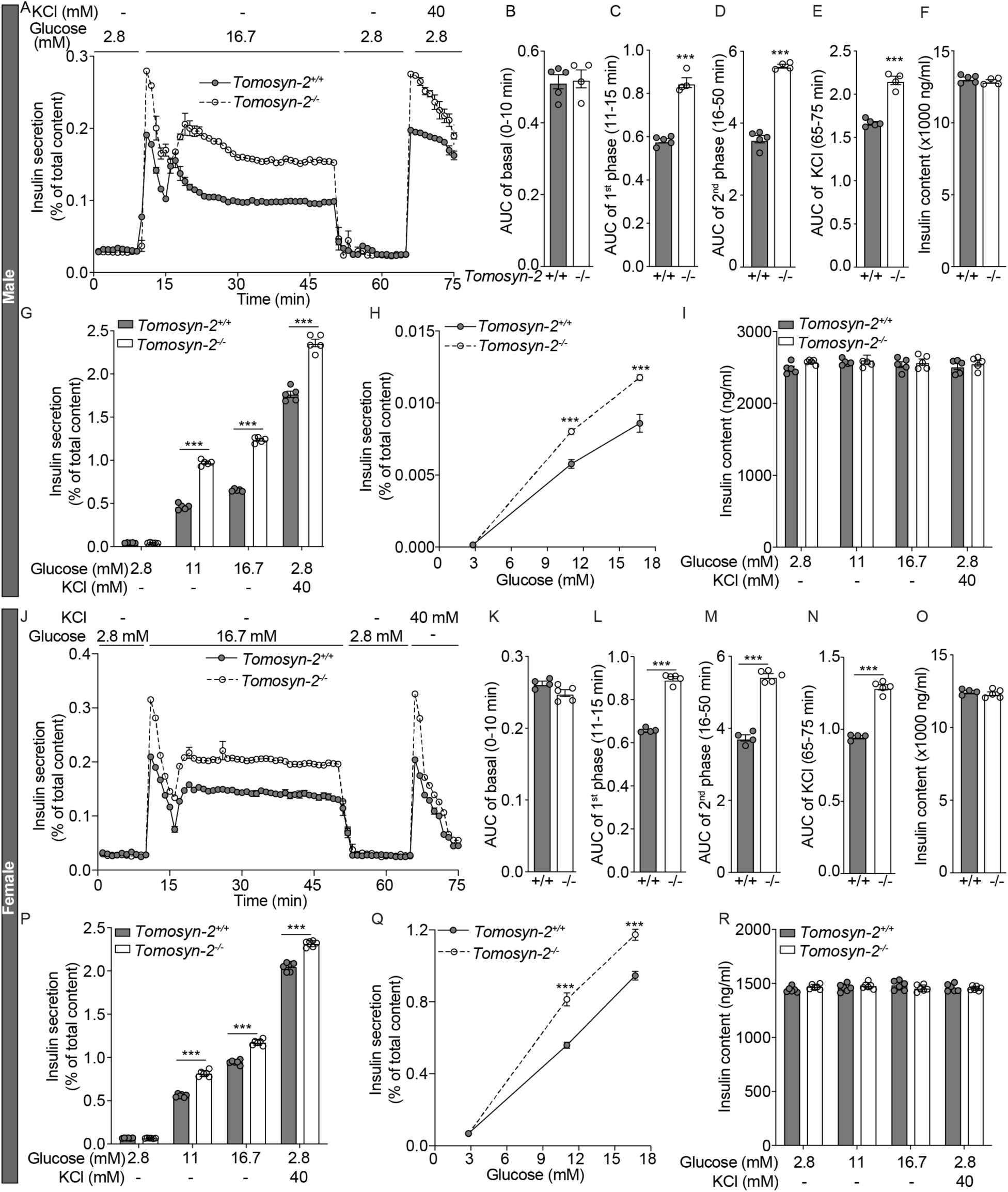
Loss of Tomosyn-2 improves biphasic insulin secretion in response to glucose from male and female mouse islets. (A) Biphasic insulin secretion from islets of *Tomosyn-2^+/+^* (closed circles, n = 5) and *Tomosyn-2^-/-^* (open circles, n = 4) 6-week-old male mice. Islets (80 per replicate) were perfused with 2.8 mM glucose, then with 16.7 mM glucose, followed by re-equilibration in 2.8 mM glucose and stimulation with 40 mM KCl. (B) AUC of basal insulin secretion (0–10 min) from (A). AUC of 1^st^ phase (11–15 min) (C) and 2^nd^ phase (16–50 min) (D) insulin secretion in response to 16.7 mM glucose from (A). (E) AUC of insulin secretion during 40 mM KCl stimulation from (A). (F) Total insulin content in islets subjected to perifusion in (A). (G) Static insulin secretion after 45 min incubation with 2.8 mM (basal), 11 mM (sub-maximal), 16.7 mM (high) glucose, and 40 mM KCl + 2.8 mM glucose in islets from *Tomosyn-2^+/+^* (closed columns) and *Tomosyn- 2^-/-^* (open columns) 6-week-old male mice (n = 5). (H) Dose-response relationship of glucose-stimulated static insulin secretion in male islets (n = 5). (J) Biphasic insulin secretion from islets of *Tomosyn-2^+/+^*(closed circles, n = 5) and *Tomosyn-2^-/-^*(open circles, n = 4) 6-week-old female mice. Islets (80 per replicate) were perfused with 2.8 mM glucose, then with 16.7 mM glucose, followed by re-equilibration in 2.8 mM glucose and stimulation with 40 mM KCl. (K) AUC of basal insulin secretion (0–10 min) from (J). AUC of 1^st^ phase (11–15 min) (L) and 2^nd^ phase (16–50 min) (M) insulin secretion in response to 16.7 mM glucose from (J). (N) AUC of insulin secretion during 40 mM KCl stimulation (65–75 min) from (J). (O) Total insulin content in islets subjected to perifusion in (J). (P) Static insulin secretion after 45 min incubation with 2.8 mM, 11 mM, 16.7 mM glucose, and 40 mM KCl + 2.8 mM glucose in islets from *Tomosyn-2^+/+^* (closed columns) and *Tomosyn-2^-/-^* (open columns) 6-week-old female mice (n = 6). (Q) Dose-response relationship of glucose-stimulated static insulin secretion in female islets (n = 5). Data are presented as mean ± s.e.m. An unpaired two-tailed Student’s t-test determined statistical significance. ^∗^P < 0.05, ^∗∗^P < 0.01, ^∗∗∗^P < 0.001.

As expected, high glucose elicited a biphasic insulin secretory response in islets from all groups. However, *Tomosyn-2^-/-^* islets secreted significantly more insulin than controls during both the early (11–16 min) and sustained (17–50 min) phases of glucose stimulation. In male islets, the early and sustained phases were increased by 37% and 44%, respectively (Fig. 3C, 3D), while female *Tomosyn-2^-/-^* islets showed 30% and 37% increases (Fig. 3K, 3L), relative to *Tomosyn-2^+/+^* controls.

KCl-induced membrane depolarization also triggered insulin secretion in both genotypes and sexes. However, *Tomosyn-2^-/-^* islets exhibited a significantly enhanced KCl-stimulated insulin secretion response, with AUCs increased by 30% in males and 44% in females (Fig. 3E, 3N). Notably, basal insulin secretion and total islet insulin content (as islets were size-matched) were unchanged between genotypes (Fig. 3D, 3F, 3I, 3K, 3O, 3R), indicating that Tomosyn-2 deletion does not affect basal insulin release under low-glucose conditions.

To further investigate β-cell secretory function, we conducted static insulin secretion assays under basal (2.8 mM), submaximal (11 mM), and maximal (16.7 mM) glucose concentrations, as well as KCl stimulation under basal glucose conditions. In both sexes, Tomosyn-2 deletion did not alter basal insulin secretion (Fig. 3G, 3P). However, insulin release in response to stimulatory glucose was markedly enhanced. Male *Tomosyn-2^-/-^* islets showed ∼2-fold increases in insulin secretion at both sub-maximal and maximal glucose concentrations (p < 0.0001; Fig. 3G, 3H), while female knockout islets exhibited ∼50% increases (p < 0.0001; Fig. 3P, 3Q). KCl-stimulated insulin secretion was also elevated by ∼50% in males and ∼20% in females (p < 0.0001; Fig. 3G, 3H, 3P, 3Q), consistent with the perifusion findings. Moreover, Tomosyn-2 sets the threshold for β-cell responsiveness to glucose and modulates sensitivity and maximal insulin secretory output. These data demonstrate that Tomosyn-2 negatively regulates insulin secretion by modulating β-cell responsiveness to both nutrient and non-nutrient stimuli, acting through mechanisms downstream of membrane depolarization.

### Tomosyn-2 inhibits SNARE complex formation via interaction with Syntaxin1A

To elucidate the molecular mechanism by which Tomosyn-2 regulates insulin secretion, we examined its interaction with Syntaxin-1A (Stx1A), a core component of the SNARE complex essential for insulin granule exocytosis. Using the Proximity Ligation Assay (PLA), which enables visualization of protein-protein interactions in situ at single-molecule resolution, in INS1 (832/13) rat β-cells. We visualized endogenous Tomosyn-2: Stx1A interactions. PLA signals, detected as red puncta, co-localized with insulin-positive (green) β-cells, confirming β-cell-specific protein interactions (Fig. 4A). Minimal signal in the control validated assay specificity. Quantitative analysis revealed a significant ∼8-fold increase in Tomosyn-2: Stx1A binding (p < 0.001; Fig. 4B). To validate this interaction independently, we performed co-immunoprecipitation (co-IP) using a stabilized V5-tagged Tomosyn-2^48^. Immunoprecipitation with anti-V5 antibody selectively pulled down both Tomosyn-2 and endogenous Stx1A, confirming Tomosyn-2: Stx1A complex formation (Fig. 4C). No signal was observed in control IPs using an isotype-matched IgG, supporting the specificity of the interaction.

**Figure 4.**
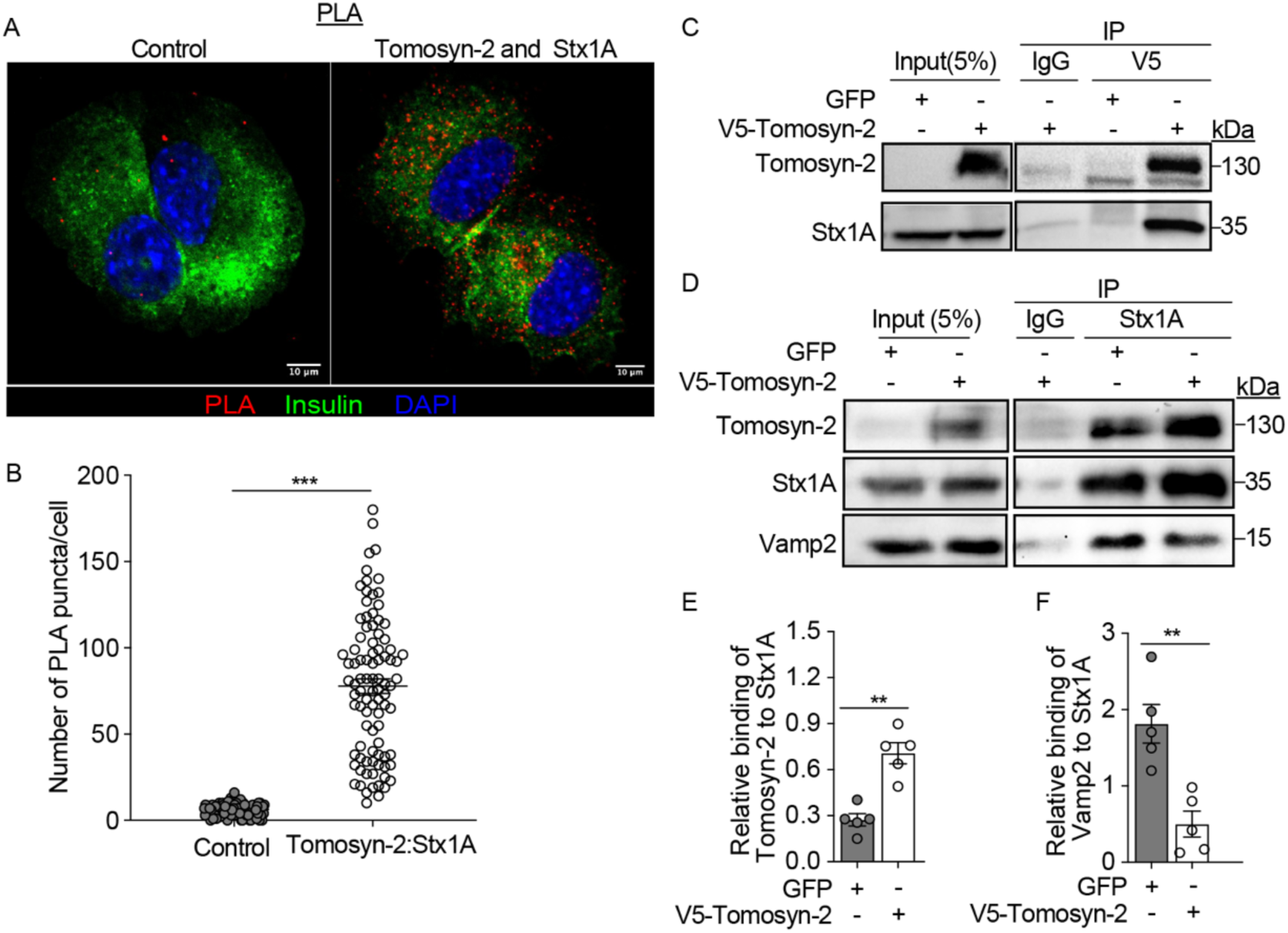
Tomosyn-2 interacts with Stx1A and reduces Stx1A–Vamp2 SNARE complex formation. (A–B) Proximity ligation assay (PLA) detecting Tomosyn-2–Stx1A interactions (red puncta) in INS1 (832/13) cells. Insulin (green) and DAPI (blue) staining mark β-cells and nuclei, respectively. (C) Quantification of Tomosyn-2–Stx1A interactions based on red PLA puncta per cell. More than 90 cells were analyzed across three independent experiments. (D) The representative Western blot shows V5-Tomosyn-2 IP results. INS1(832/13) β-cells were transfected with V5-Tomosyn-2 or V5-GFP plasmids for 48 h. IP was performed using an anti-V5 antibody in lysis buffer containing either 1 mM Ca²⁺ (left) or 2 mM EGTA (right) (n = 3). (E) The representative Western blot showing Stx1A IP followed by the co-IP of Vamp2 and Tomosyn-2. (F–G) Quantification of immunoblot band intensity using ImageJ, showing the relative binding of Tomosyn-2 (F) or Vamp2 (G) to Stx1A (n = 5). Data are presented as mean ± s.e.m. An unpaired two-tailed Student’s t-test determined statistical significance. ^∗^P < 0.05, ^∗∗^P < 0.01, ^∗∗∗^P < 0.001.

We next examined whether Tomosyn-2 affects SNARE complex formation. Tomosyn-2 was transiently overexpressed in INS1(832/13) cells, and Stx1A was immunoprecipitated using anti-Stx1A antibody and analyzed for associated SNARE proteins. Tomosyn-2 resulted in a ∼3-fold increase in Tomosyn-2-bound to Stx1A (p < 0.001) compared to control cells (Fig. 4D–E). Strikingly, the association of Vamp2 with Stx1A was reduced by approximately 75% (p < 0.001) in the presence of Tomosyn-2, suggesting that Tomosyn-2 attenuates Vamp2 from the Stx1A-SNARE complex (Fig. 4F).

Together, these results demonstrate that Tomosyn-2 directly interacts with Stx1A in β-cells and antagonizes SNARE complex assembly by reducing Stx1A–Vamp2 interactions. These findings reveal a molecular mechanism by which Tomosyn-2 functions as a negative regulator of insulin secretion, inhibiting SNARE-mediated granule fusion.

### Tomosyn-2 reciprocally modulates the expression of genes associated with insulin secretion and proliferation

During postnatal development, pancreatic β-cells undergo transcriptional and functional maturation, transitioning from a proliferative state to one characterized by robust, stimulus-coupled insulin secretion (Fig. 1A–N). Consistent with this trajectory, our data show that the loss of Tomosyn-2 increases insulin secretion, thereby improving glucose clearance in 6-week-old mice (Fig. 2I–L). To investigate how Tomosyn-2 influences the molecular events governing β-cell maturation, we performed RNA-seq analysis on islets isolated from *Tomosyn-2^-/-^* and *Tomosyn-2^+/+^* male mice.

Differential expression analysis identified 4,784 genes significantly altered by Tomosyn-2 deletion (Fig. 5A, Supplementary Table 4), with 18.7% were upregulated and 13.9% were downregulated in *Tomosyn-2^-/-^* islets (Fig. 5B). Upregulated transcripts included key regulators of β-cell identity and insulin secretion, such as *Ins1*, *Ins2*, *NeuroD1*, *Mdm1*, *Pdx1 Cxcl13*, and *Xbp1*. In contrast, genes associated with cell proliferation— including *Sox1, Akt1, Pik2cd, Ccna1, Foxm1, Pcna,* and *Mki67*—were significantly downregulated.

**Figure 5.**
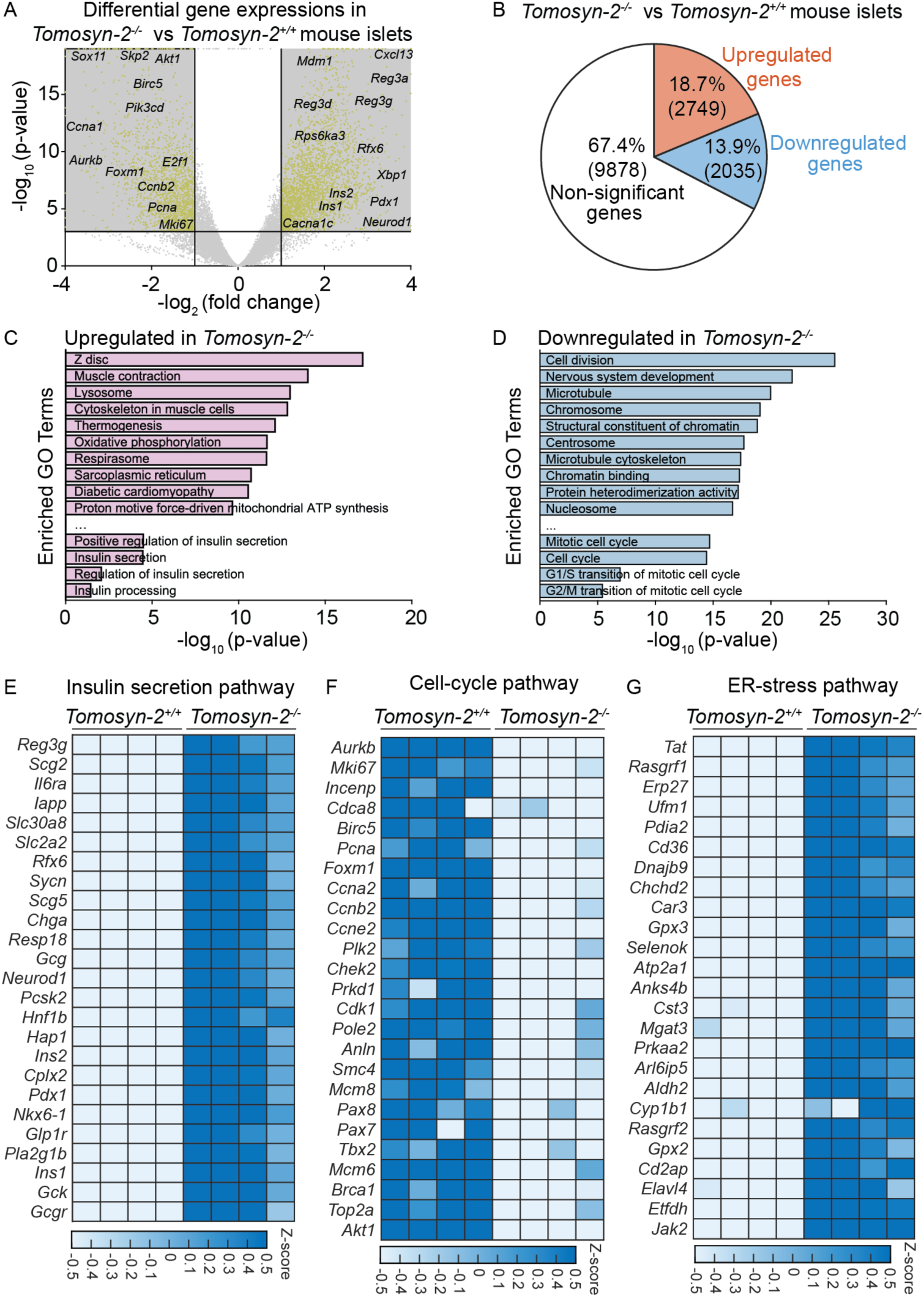
Bulk RNA-seq analysis reveals that loss of Tomosyn-2 reciprocally regulates gene networks associated with insulin secretion and β-cell proliferation. (A) The association between the base-2 logarithm of fold-change and the base-10 logarithm of p-value, computed by the Deseq2 pipeline, was visualized using a volcano plot; significantly differentially expressed genes were plotted as yellow dots (n = 4 per group). B) The ratio of genes upregulated, downregulated genes, and genes that were neither up-nor down-regulated in *Tomosyn-2^-/-^* vs. *Tomosyn-2^+/+^* islets was visualized in a pie chart. C-D) Top 10 enriched GO terms and KEGG pathways, insulin-secretion, and cell-cycle were visualized in a bar chart; the x-axis represents the base-10 logarithm of p-value for each term/pathway. E-G) The abundance of expression (normalized via z-score) for insulin-secretion (E), cell-cycle (F), and ER-stress (G) genes in each cluster is displayed as a heat map.

Gene Ontology (GO) and KEGG pathway enrichment analyses revealed 364 pathways enriched among upregulated genes in *Tomosyn-2^-/-^* islets, including those involved in insulin secretion, β-cell identity, insulin processing, and positive regulation of insulin secretion (Fig. 5C; Supplementary Table 5). Conversely, 751 pathways were enriched among downregulated genes, predominantly associated with mitotic progression, cell cycle control, and G1/S and G2/M transitions (Fig. 5D). These data suggest that Tomosyn-2 suppresses β-cell secretory capacity while promoting proliferative programs during postnatal development.

A closer inspection of insulin secretion–related genes revealed upregulation of transcriptional and functional regulators of insulin synthesis and release, including *Iapp, Slc2a2, Syncn, Scg5, Chga, Neurod1, Ins2, Pdx1, Nkx6.1, Glp1r, Ins1, Gck, and Gcgr* due to the loss of Tomosyn-2 (Fig. 5E). In contrast, proliferation-associated transcripts— including *Aurkb*, *Mki67*, *Ccna2*, *Ccne2*, *Akt1*, *Brca1*, and *Ezh2*—were consistently downregulated (Fig. 5F). Notably, genes associated with endoplasmic reticulum (ER) stress and adaptive unfolded protein response, including *Xbp1*, *Tat*, *Erp27*, *Gpx3*, *Cd36*, and *Jak2*, were also upregulated (Fig. 5G), indicating a potential adaptive response accompanying heightened insulin biosynthetic demand.

Heatmap and hierarchical clustering analyses further supported the reciprocal regulation of gene networks promoting β-cell functional identity versus proliferation (Supplementary Fig. 5). Network analysis (via STRING v.12^62^ database) showed that differentially expressed genes associated with insulin secretion strongly interacted, as well as genes of ER-stress and cell-cycle pathways (Fig. 6A-C). Network analysis of upregulated genes revealed a densely connected cluster centered on key β-cell regulators including *Ins1*, *Ins2*, *Kcnj11, Abcc8, Slc2a2, NeuroD1*, *Glp1r*, and *Ucn3* (Fig. 6A). A parallel network analysis of upregulated ER stress–related genes identified *Xbp1*, *Ern1*, *Bcl2l1*, and *Map3k5* as central hub genes (Fig. 6B). Downregulated genes formed a tightly connected cell cycle module, anchored by *Akt1*, *Ezh2*, *Foxm1, Prr1, Cep55*, *Aurkb*, *Plk1*, and *Trip13* as central nodes (Fig. 6C).

**Figure 6.**
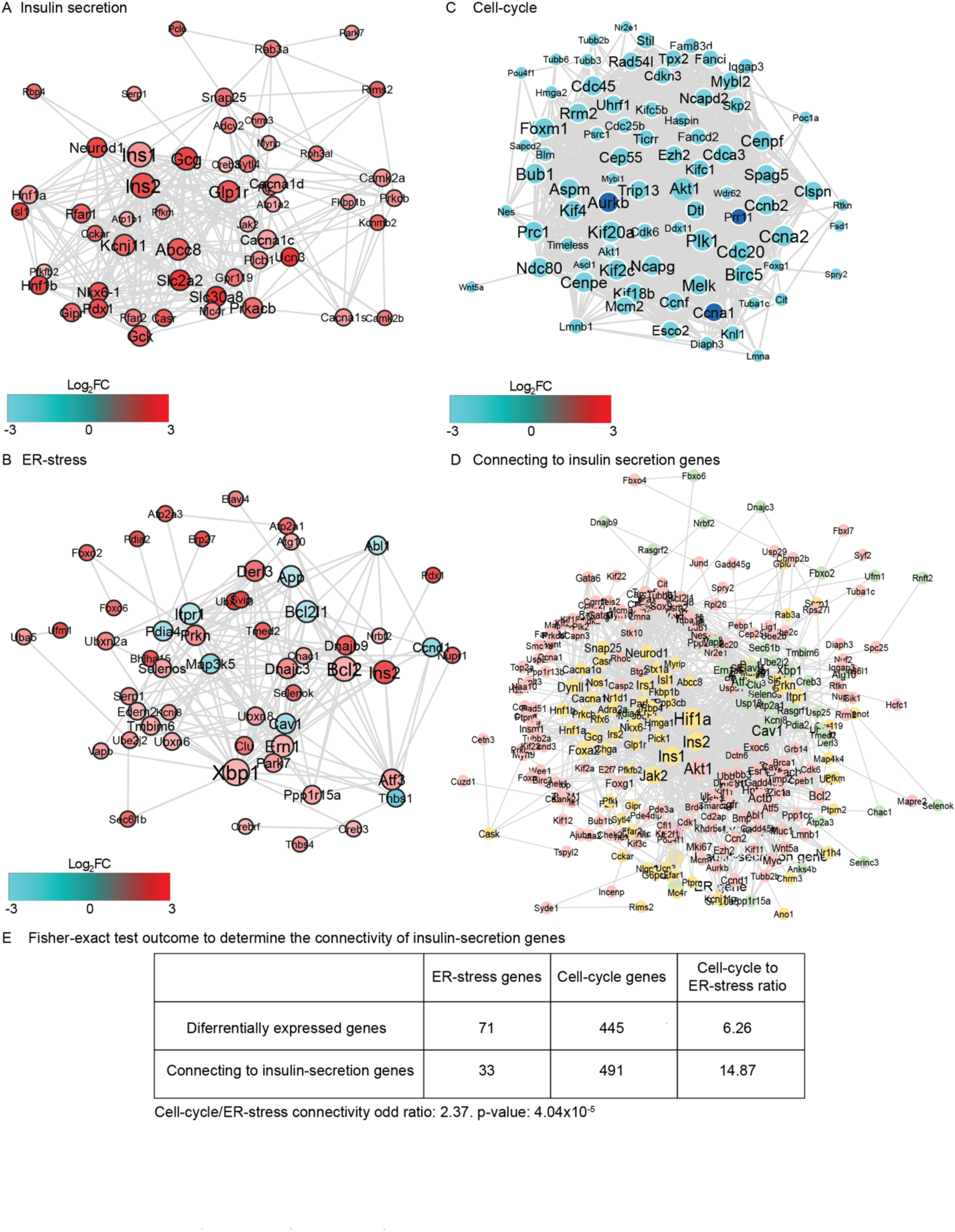
Intersecting network analysis reveals interactions among genes involved in insulin secretion, cell cycle, and ER stress pathways. (A-C) Network among differentially-expressed gene of insulin-secretion (A), ER-stress (B), and cell-cycle (C); the networks were queried via STRING v.12 database and visualized via Cytoscape software; each node represent a gene, the node size represents the number of interaction for each gene; and the node color represent the base-2 logarithm of fold-change expression between *Tomosyn-2^+/+^* and *Tomosyn-2^-/-^* islets. D) Crosstalk network among insulin-secretion, ER-stress, and cell-cycle genes; the node size represents the number of interactions for each gene. E) KEGG Pi3k-Akt signaling pathway was visualized with color-code representing base-2 logarithm of fold-change expression between *Tomosyn- 2^+/+^* and *Tomosyn-2^-/-^*islets; red: upregulated in *Tomosyn-2^-/-^*, cyan: downregulated in *Tomosyn-2^-/-^*, green: neither up-nor down-regulated.

To assess potential crosstalk among these pathways, we performed Fisher’s exact test to evaluate DEG overlap. Of 71 ER stress–associated DEGs, 33 showed connectivity with insulin secretion genes, while 445 downregulated cell cycle genes were linked to 491 upregulated insulin secretion genes. The odds ratio for connectivity between the cell cycle and insulin secretion networks was 2.37 (p = 4.04 × 10^^5^), indicating a strong reciprocal relationship. Notably, insulin secretion genes were over twice as likely to interact with cell cycle genes than with ER stress–related genes (Fig. 6D). Among the shared nodes in this tripartite network, *Ins1*, *Ins2*, *Jak2*, *Hif1a*, and *Akt1* emerged as key integrators and were also components of the KEGG PI3K-Akt signaling pathway (Fig. 6D–E).

Collectively, these findings suggest that Tomosyn-2 acts as a molecular brake on insulin secretion during early postnatal development, potentially dampening insulin secretion– associated gene networks while promoting cell cycle progression to facilitate β-cell proliferation. However, its loss drives a gene expression program favoring β-cell functional competence at the expense of proliferative capacity.

### Loss of Tomosyn-2 decreases β-cell proliferation via the AKT1 signaling pathway

Our network analysis identified **Akt1** as a central hub gene linking insulin secretion and cell cycle regulatory networks in response to Tomosyn-2 deletion (Fig. 6D). Given the well-established role of the PI3K-Akt1-Cyclin D1 axis in promoting cell cycle progression and β-cell proliferation, we examined whether Tomosyn-2 modulates this pathway (KEGG pathway: mmu04151; https://www.kegg.jp/pathway/mmu04151). Genes in the pathway were color-coded based on RNA-seq differential expression data comparing *Tomosyn-2^-/-^* and *Tomosyn-2^+/+^* mouse islets: red for upregulated, cyan for downregulated, and green for unchanged genes (Supplementary Fig. 4). Based on this visualization, it appears that the loss of Tomosyn-2 decreased the expression of Irs1– PI3K–Akt1–Ccnd1 signaling module, even as the expression of Ins1/2 and insulin receptor was elevated. Suggesting that the loss of Tomosyn-2 regulates proliferation by reducing the PI3K-Akt1 signaling pathway. Thus, we evaluated the effect of Tomosyn-2 on the PI3K-Akt1 signaling pathway, which is central to β-cell proliferation^63^.

RNA-seq analysis revealed significant downregulation of multiple components of the PI3K-Akt1 pathway, including *Irs1*, *Irs2*, *Pik3cd*, and *Akt1*, along with cell cycle regulators *Ccnd1*, *Ccnd3*, *Ccne2*, *Myc*, *Cdk2*, and *Cdk6* in *Tomosyn-2^-/-^*islets (Fig. 7A). These data implicate reduced PI3K-Akt1 signaling and impaired cell cycle progression in the absence of Tomosyn-2. Consistent with transcriptomic findings, Western blot analysis showed a marked reduction in phosphorylated (p)-Akt1 protein levels in *Tomosyn-2^-/-^* islets, as well as a trend in reduction total(t)-Akt1 (Fig. 7B, quantified in Fig. 7C). Given that Akt1 phosphorylation enhances *Cyclin D1* expression to promote β-cell proliferation^64^, we assessed Cyclin D1 protein abundance and found a 75% reduction in Cyclin D1 levels in *Tomosyn-2^-/-^*islets (Fig. 7B, quantified in Fig. 7D). In contrast, the protein level of the cell cycle inhibitor *Cdkn1b* (*p27Kip1*) was elevated nearly threefold (Fig. 7B, quantified in Fig. 7E), further supporting a shift toward cell cycle arrest.

**Figure 7.**
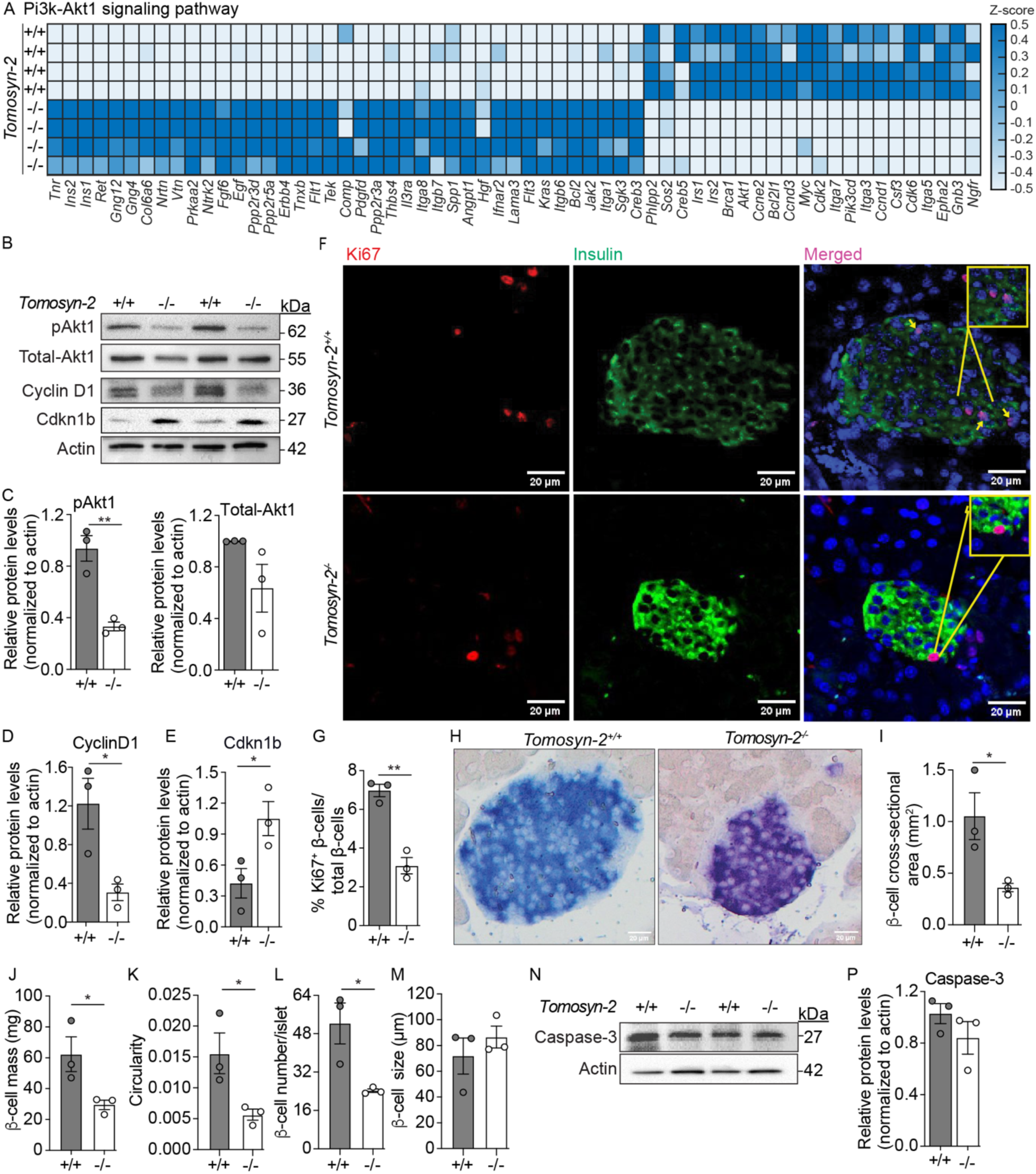
Loss of Tomosyn-2 reduces β-cell proliferation by downregulating Akt signaling, without affecting apoptosis. (A) Heatmap showing significantly differentially expressed genes associated with the PI3K–Akt1 signaling pathway in islets of *Tomosyn- 2^-/-^* mice compared to *Tomosyn-2^+/+^* mice. (B) Representative Western blot showing protein levels of total Akt1, phosphorylated Akt1 (pAkt1), Cyclin D1, and Cdkn1b in islets (n = 3). (C–E) Quantification of Western blot band intensities using ImageJ for (C) total Akt, (D) p-Akt, and (E) Cyclin D1, normalized to β-actin. (F) IF staining of islets from *Tomosyn-2^+/+^* and *Tomosyn-2^-/-^* mice shows Ki67 (red), insulin (green), and DAPI-stained nuclei (blue). Images captured by confocal microscopy at 20X magnification. Arrows indicate Ki67 insulin^+^ β-cells; insets show magnified views. (G) Quantification of proliferating β-cells (% Ki67^+^ insulin^+^β-cells per total insulin+β-cells) using ImageJ (N = 3). (H) Insulin-positive islets were visualized by the presence of blue chromogenic staining in *Tomosyn-2^-/-^* and *Tomosyn-2^+/+^* mouse pancreata. Quantification of β-cell cross-sectional area (I), circularity (J), β-cell count per islet (K), β-cell number/islet (L), and β-cell size (M). (N) The representative Western blot shows levels of apoptosis marker Caspase-3 in islets (n = 3). (M) Quantification of activated Caspase-3 protein levels, normalized to β-actin. Data are presented as mean ± SEM. An unpaired two-tailed Student’s t-test determined statistical significance. *P < 0.05, **P < 0.01, ***P < 0.001.

To evaluate the functional consequences of altered Akt1 signaling on β-cell proliferation, we performed IF analysis on neonatal pancreata, co-staining for Ki67 (a marker of proliferation) and insulin. *Tomosyn-2^-/-^* islets showed a significant reduction— approximately 50%—in Ki67^+^ insulin+β-cells compared to controls (Fig. 7F–G). A similar decline in Ki67^+^ non–β-cells was also observed (Supplementary Fig. 5), suggesting that Tomosyn-2 loss broadly suppresses islet cell proliferation. Morphometric analyses further revealed that Tomosyn-2 deletion leads to a reduction in β-cell mass, cross-sectional area, circularity, and β-cell number (Fig. 7H–K). However, no significant change in average β-cell size was detected. These results suggest that loss of Tomosyn-2 compromises β-cell mass by impeding proliferation.

Because *Tomosyn-2^-/-^* islets exhibited upregulation of genes involved in ER stress (Fig. 5G), we considered whether apoptosis might contribute to the observed reduction in β-cell mass. However, neither qPCR nor Western blot analysis revealed increased levels of cleaved caspase-3 (Fig. 7M, quantified in Fig. 7N), indicating that ER stress does not lead to apoptosis in this context. This suggests that β-cells may engage in adaptive unfolded protein response (UPR) pathway to buffer against ER stress (Supplementary Fig. 6) and preserve cell viability despite the loss of Tomosyn-2.

Together, these findings demonstrate that Tomosyn-2 supports β-cell proliferative capacity during postnatal development by sustaining PI3K-Akt1 pathway signaling. Its deletion leads to impaired Akt1 phosphorylation, reduced Cyclin D1 abundance, and increased p27Kip1 expression, thereby suppressing cell cycle progression and limiting β-cell mass expansion without inducing apoptosis.

### Loss of Tomosyn-2 enhances β-cell identity and functional maturation

To assess the role of Tomosyn-2 in regulating β-cell identity and maturation, we examined the expression of canonical markers in islets isolated from *Tomosyn-2^-/-^*and *Tomosyn- 2^+/+^* mice. *Tomosyn-2^-/-^* islets exhibited significantly elevated mRNA abundance of key β-cell identity transcription factors, including *Nkx6.1*, *NeuroD1*, *Pdx1*, *Hnf4α*, *Isl1*, and *Foxo1*, as well as markers of β-cell functional maturity such as *Ucn3*, *Ins1*, and *Ins2* (Fig. 8A). In contrast, expression of genes associated with immature β-cell states—including *Ldha*, *Hk1*, *Hk2*, and *Rest*—was significantly downregulated. Importantly, there were no significant changes in the expression of β-cell dedifferentiation markers (Fig. 8B), suggesting that Tomosyn-2 deletion promotes β-cell maturation without inducing dedifferentiation.

**Figure 8.**
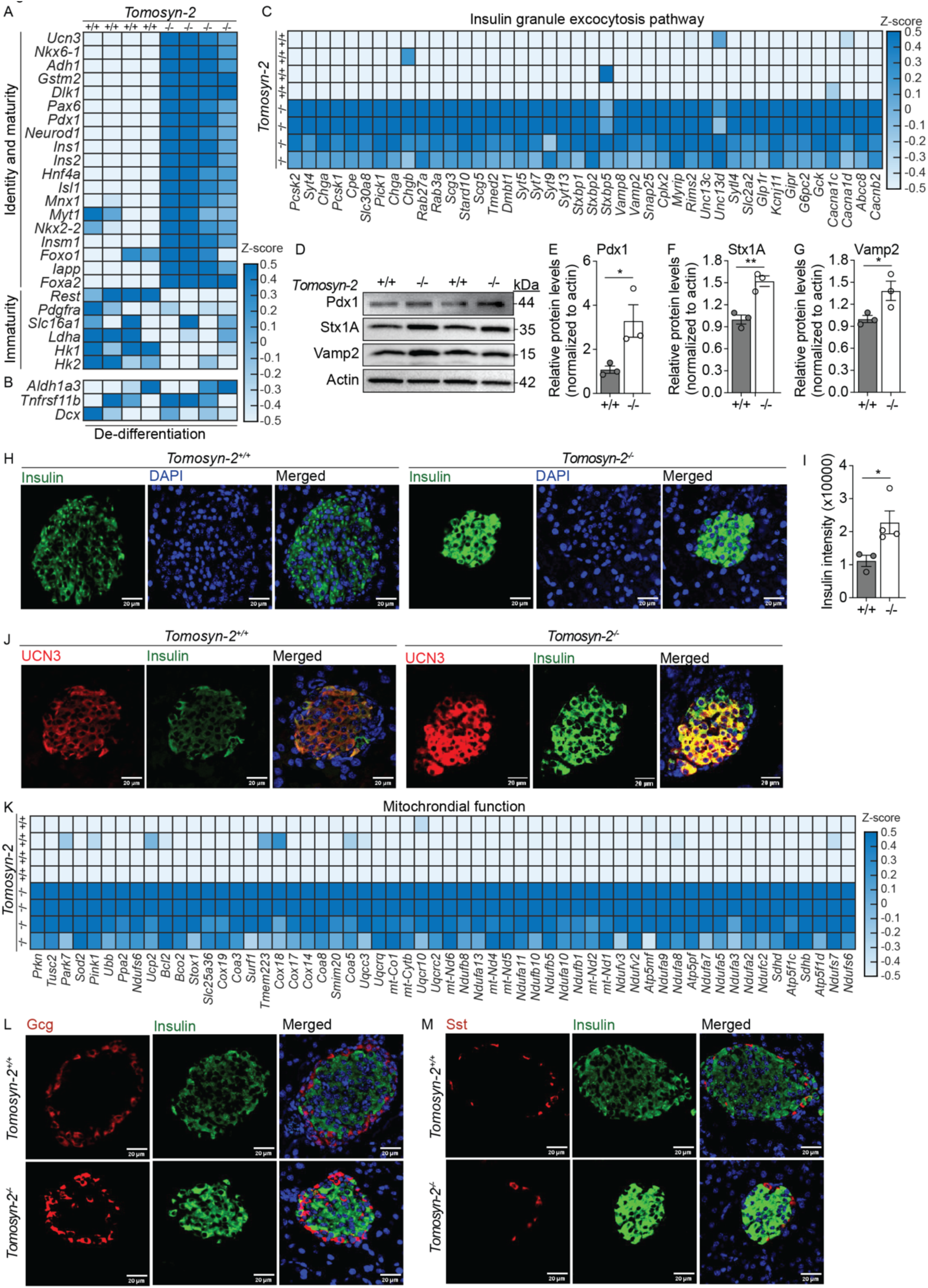
Loss of Tomosyn-2 increases β-cell identity and functional maturity. (A– C) Heatmap representation of significantly differentially expressed genes regulating (A) β-cell maturity and immaturity, (B) de-differentiation, and (C) insulin granule exocytosis in islets of *Tomosyn^-^2^+/+^* and *Tomosyn-2^-/-^* mice. (D) Representative image of Western blot analysis of proteins regulating β-cell maturity (Pdx1) and exocytosis (Stx1A, Vamp2) (n = 3). (E–G) Quantification of protein band intensities using ImageJ, assessing the relative abundance of (E) Pdx1, (F) Stx1A, and (G) Vamp2 normalized to actin. (H) Immunostaining showing the expression of insulin (green) and DAPI (blue) in islets of *Tomosyn^-^2^+/+^* and *Tomosyn-2^-/-^*mice (n = 3). (J) Quantification of insulin intensity by ImageJ assessing insulin expression in β-cells of *Tomosyn^-^2^+/+^* and *Tomosyn-2^-/-^* mice (n = 3). (K) Immunostaining showing the expression of β-cell maturation marker Ucn3 (red) and insulin (green) in islets of *Tomosyn^-^2^+/+^* mice compared to *Tomosyn-2^-/-^*(n = 3). (L) Heatmap representation of significantly differentially expressed genes regulating β-cell mitochondrial metabolism in *Tomosyn^-^2^+/+^* mice compared to *Tomosyn-2^-/-^*(n = 4). (M) Immunostaining showing expression of α-cell marker glucagon (red) and β-cell marker insulin (green) in islets of *Tomosyn^-^2^+/+^*mice compared to *Tomosyn-2^-/-^* (n = 3). (N) Immunostaining showing expression of δ-cell marker somatostatin (red) and β-cell marker insulin (green) in islets of *Tomosyn^-^2^+/+^*mice compared to *Tomosyn-2^-/-^* (n = 3). All immunostaining images were acquired using a confocal microscope at 20× magnification. An unpaired two-tailed Student’s t-test determined statistical significance. Data are presented as mean ± SEMs. P < 0.05, ^∗∗^P < 0.01, ^∗∗∗^P < 0.001.

Functionally mature β-cells are defined by their capacity to synthesize, process, and secrete insulin. Consistent with this, *Tomosyn-2^-/-^* islets showed increased expression of genes involved in proinsulin processing (*Pcsk1*, *Pcsk2*, *Cpe*, *Chga*, *Chgb*, *Scg5*), GSIS signaling (*Gck*, *Glp1r*, *Gipr*, *Slc2a2*, *Kcnj11*, *Cacna1*), and insulin granule docking and fusion (*Rab3a*, *Rab27a*, *Syt7*, *Syt5*, *Syt4*, *Vamp2*, *Vamp8*, *Stxbp2*, *Snap25*, *Stxbp5*, *Unc13c*, and *Rims2*) (Fig. 8C).

Western blot analysis confirmed increased protein abundance of the β-cell identity factor Pdx1 (3-fold, *p* = 0.042), and exocytosis-related proteins Stx1A (1.5-fold, *p* = 0.005) and Vamp2 (1.4-fold, *p* = 0.051) in *Tomosyn-2^-/-^* islets compared to controls (Fig. 8D; quantified in Fig. 8E–G). IF staining of pancreatic sections revealed a significant increase in insulin protein levels (2-fold, *p* = 0.04) in *Tomosyn-2^-/-^* β-cells (Fig. 8H–I). Similarly, Ucn3—a hallmark of mature β-cells—was markedly increased in insulin+β-cells of *Tomosyn-2^-/-^*islets (Fig. 8J).

Given the association between β-cell maturity and enhanced mitochondrial metabolism, we analyzed the expression of mitochondrial genes. A broad upregulation of mitochondrial transcripts was observed in *Tomosyn-2^-/-^* islets (Fig. 8K), further supporting a shift toward a mature, metabolically active β-cell state.

Together, these data indicate that Tomosyn-2 negatively regulates the acquisition of β-cell identity and functional maturity during postnatal development. Its loss activates a gene expression program that enhances insulin production, granule exocytosis, and mitochondrial function, hallmarks of a mature and functional β-cell phenotype.

### Loss of Tomosyn-2 alters islet architecture without disrupting β-cell identity

In rodent islets, β-cells are typically localized to the core, surrounded by α- and δ-cells at the periphery—an organization essential for optimal intercellular communication and β-cell functional maturation in adulthood. To determine whether Tomosyn-2 influences islet cytoarchitecture, we performed immunostaining to assess the spatial distribution of α-, β-, and δ-cells in *Tomosyn-2^-/-^* and *Tomosyn-2^+/+^* pancreata. *Tomosyn-2^-/-^* islets exhibited an altered distribution of α-cells, which were aberrantly positioned in both the peripheral and core regions of the islets (Fig. 8L), deviating from the canonical peripheral localization observed in controls. In contrast, the spatial organization of δ-cells remained unchanged (Fig. 8N). Notably, Tomosyn-2 deletion did not alter the expression of β-cell dedifferentiation markers; instead, it enhanced β-cell identity and maturation signatures, suggesting that core infiltration by α-cells is unlikely to result from β-cell loss of identity or dedifferentiation. Rather, the disrupted islet architecture may be attributed to the reduced β-cell proliferation observed in *Tomosyn-2^-/-^* islets, potentially leading to insufficient β-cell mass to maintain normal core occupancy. Further investigation is required to elucidate the mechanisms underlying α-cell mislocalization in the absence of Tomosyn-2, particularly in the context of preserved β-cell identity. Future studies will aim to define the cellular and molecular cues responsible for this altered spatial organization.

## Discussion

The mechanisms governing the balance between proliferative, immature β-cells and their transition to functionally mature β-cells during postnatal development remain poorly understood. In this study, we show that biphasic glucose-stimulated insulin secretion (GSIS) increases with age, inversely correlating with a progressive decline in β-cell proliferation. This maturation process coincides with a marked reduction in Tomosyn-2 protein abundance—an endogenous inhibitor of insulin secretion—suggesting a functional role for Tomosyn-2 in regulating β-cell maturation. Tomosyn-2-deficient mice exhibit improved glucose tolerance due to enhanced insulin secretion, driven by reduced Tomosyn-2 binding to Stx1A. This reduction enables decreased assembly of Stx1A-containing SNARE complexes, which are essential for insulin granule fusion with the plasma membrane. RNA-seq network analyses reveal strong inverse connectivity between genes involved in insulin secretion and those associated with cell proliferation, supporting the functional data. Mechanistically, the loss of Tomosyn-2 suppresses Akt1 signaling, leading to reduced β-cell proliferation and a decrease in β-cell mass. Simultaneously, β-cells exhibit enhanced identity and functional maturity, with diminished expression of immaturity markers and altered islet cytoarchitecture. Together, these findings establish a cytosolic factor, Tomosyn-2, which reciprocally regulates insulin secretion and β-cell proliferation, thereby influencing β-cell maturation and mass during the postnatal period. *Thereby, identifying Tomosyn-2 as a dual regulator of insulin secretion and β-cell proliferation in the postnatal development*.

While reduced insulin secretion contributes to hyperglycemia in insulin-resistant states^65–67^, chronic β-cell overstimulation may also lead to hyperinsulinemia and eventual β-cell exhaustion, contributing to the development of type 2 diabetes (T2D)^68,69^. These prevailing models highlight that both insufficient and excessive insulin secretion can underlie β-cell dysfunction in T2D. In our study, Tomosyn-2 deletion reduces β-cell proliferation and mass while enhancing ex vivo insulin secretion, increasing plasma insulin levels, and improving glucose clearance. These effects occur without changes in body weight or insulin sensitivity in young mice, suggesting a β-cell-intrinsic mechanism independent of systemic insulin resistance. It remains possible that increased insulin secretion may exert autocrine or paracrine effects on β-cells, potentially leading to desensitization of insulin receptor signaling, attenuated Akt1 signaling, and consequently, proliferation. However, the direct role of β-cell-derived insulin in modulating β-cell function remains uncertain^35,36^. Interestingly, Tomosyn-2 deficiency results in upregulation of Ins1, Ins2, and the insulin receptor (IR), while downregulating Irs1, Irs2, Pi3k, Akt1, and Ccnd1 (Fig. 7, Supplementary Fig. 4). These data suggest that suppression of the PI3K-Akt1 signaling pathway in Tomosyn-2-deficient β-cells may occur independently of insulin receptor. The precise molecular mechanism by which Tomosyn-2 regulates Akt1 signaling and β-cell proliferation remains to be elucidated. Whether the inhibition of β-cell proliferation in Tomosyn-2-deficient mice results from diminished insulin receptor signaling via an autocrine, paracrine, or intracellular β-cell signaling mechanism remains to be elucidated.

*How does Tomosyn-2 regulate insulin secretion?* Tomosyn-2 was identified as a gene located on chromosome 16 within a fasting glucose locus in an F2 mouse cross of C57BL/6J. *Lep^OB^* with BTBR. *Lep^OB^*^59^. Positional cloning revealed that a SNP in the coding region of the *Tomosyn-2* gene led to increased Tomosyn-2 protein abundance, associated with the hyperglycemia-hyperinsulinemia phenotypes in these mice, which were attributed to reduced ex vivo islet insulin secretion^59^. Similarly, a gain-of-function mutation (Val1043Ile) in the human *Tomosyn-2* gene has been shown to enhance Tomosyn-2-mediated inhibition of exocytosis^55^. Published studies show that increased abundance of Tomosyn-2 decreases insulin secretion, leading to diabetogenic phenotypes^48,59^. Herein, in elucidating the molecular mechanism of Tomosyn-2 and its physiological role, we observed that the loss of Tomosyn-2 increases insulin secretion, concomitantly decreasing β-cell mass—a trait known to enhance susceptibility to the development of T2D. These studies highlight the importance of tightly regulating Tomosyn-2 protein abundance during postnatal β-cell maturation, as alterations in Tomosyn-2 protein levels lead to β-cell dysfunction in mouse models, potentially increasing the risk of developing T2D.

The molecular mechanism by which Tomosyn-2 inhibits insulin secretion is not completely known. We demonstrate that loss of Tomosyn-2 increases both phases of insulin secretion in *Tomosyn-2^-/-^* islets compared to control *Tomosyn-2^+/+^* islets. The formation of the Stx1A-SNARE complexes regulates biphasic insulin secretion^70^. We demonstrate the endogenous binding of Tomosyn-2 with Stx1A and that Tomosyn-2 directly inhibits the formation of Stx1A-SNARE complexes (Fig. 4), suggesting the inhibitory effect of Tomosyn-2 on biphasic insulin secretion, identifying the mechanism by which Tomosyn-2 inhibits insulin secretion. Insulin granules undergo fusion to the PM from distinct cellular pools in the biphasic GSIS. Pre-docked granules are present near the PM and immediately undergo fusion upon stimulation, contributing to early-phase insulin secretion. While newcomer insulin granules are present in the cytosol, away from the PM. They undergo fusion by two distinct modes, short-dock or no-dock, contributing to both phases of GSIS. The underlying mechanism affecting the fusion of insulin granules that undergo docking (pre-docked or short-dock) vs. no-dock is not clearly understood. Stx1A facilitates the fusion of pre-docked and newcomer-short-docked insulin granules that have increased resident times at the PM^70^. As Tomosyn-2 binds to Stx1A under conditions of insulin secretion (Fig. 4A), it is plausible that Tomosyn–2–Stx1A binding imparts a temporal constraint on insulin granules at the insulin granule-PM interface, reducing the ability of a subset of insulin granules to undergo fusion. The mechanism by which Tomosyn-2 modulates insulin granule fusion modes remains to be elucidated.

*How the loss of a cytosolic protein, Tomosyn-2, decreases β-cell mass*. β-cell proliferation is crucial for postnatal expansion of the β-cell mass. Proliferating β-cells are functionally immature^71^, exhibiting reduced insulin secretion in response to glucose stimulation. At birth, β-cell proliferation rates are high but decline progressively with age, as reported here (Fig. 1) and by others, reaching 1-2% in adult mice. Therefore, it is necessary to understand the mechanisms that regulate the balance between proliferating and insulin-secreting β-cells to establish a threshold of functional β-cell mass, which is key to maintaining whole-body glucose homeostasis in adults.

Insulin receptor signaling is known to regulate β-cell proliferation^72^, where the activation of the insulin receptor substrate-2 (Irs2)-Pi3k-Akt1 signaling module regulates downstream effectors, including Foxo1, Gsk3β, CyclinD1, p27Kip1 (Cdkn1b), and p21Cip1 (Cdkn1a), for cell cycle progression. We demonstrate that Tomosyn-2 loss significantly reduces the expression of Irs1, Irs2, Akt1, and Gsk3β, as well as the protein levels of phospho-Akt1 (Fig. 7). Consequently, observed is the reduced expression of gene involved in cell-cycle progression, and protein abundance of a key cell cycle regulator, CyclinD1, and Ki67^+^β-cells, along with increased levels of cell cycle inhibitor Cdkn1b. These outcomes demonstrate that Tomosyn-2 deficiency reduces Akt1 signaling, thereby decreasing β-cell proliferation. The mechanism by which Tomosyn-2 loss regulates the Akt1-mediated reduction in β-cell proliferation remains to be determined. Whether Tomosyn-2 directly binds components of the insulin signaling pathway or acts through secondary mechanisms, such as elevated insulin production, as reduced insulin production is known to alter gene expression profiles known to increase β-cell proliferation^5,73^, autocrine insulin signaling, or altered gene expression, remains an open question. The observed downregulation of Creb5 and Irs2—key regulators of β-cell mass^74,75^ —further supports this possibility. Determining the mechanism by which Tomosyn-2 regulates Akt1 signaling and cell cycle progression remains an area for future investigation. This study shows that Tomosyn-2 regulates insulin secretion by modulating the formation of Stx1A-SNARE complex and also reduces β-cell mass by decreasing Akt1-mediated β-cell proliferation. Thereby, highlighting the reciprocal control by Tomosyn-2 on insulin secretion and proliferation, two key cellular processes in postnatal β-cells.

Tomosyn-2 deficiency reduces β-cell number, circularity, and cross-sectional area, leading to smaller islets with intermingled α-cells and disrupted core-mantle architecture. Interestingly, this architectural disorganization does not reflect β-cell dedifferentiation. On the contrary, markers of β-cell identity (*Pdx1, Nkx6.1, NeuroD1*) and maturity (Ins1, Ins2, Ucn3, mitochondrial oxidative phosphorylation genes, insulin granule trafficking, and exocytosis genes) are increased, while immaturity and disallowed genes (*Ldha, Rest, Pdgfra, Slc16a1, Hk1, Hk2*) are suppressed. This suggests that the observed architectural changes may result from reduced β-cell proliferation rather than loss of identity. Islet architecture, which arises from differential growth and sorting of α- and β-cells, may thus be altered by disproportionate proliferation between the two cell types.

Tomosyn-2 is an inhibitor of insulin secretion^48,59^ (Figs. 2, 3). Tomosyn-2 is highly expressed in islets between postnatal days 7–14, with levels declining into adulthood. We propose that this temporal pattern restricts early insulin secretion, enabling sufficient β-cell proliferation before allowing functional maturation. Our data support this model: Tomosyn-2 loss increases insulin secretion accelerates β-cell identity and functional maturity evident by increased expression of identity (*Nkx6.1, NeuroD1, Pdx1*) and maturity markers (*Ucn3, Ins1, and Ins2*), mitochondrial oxidative phosphorylation, insulin granule trafficking, and insulin exocytosis along with the reduction of immaturity markers (*Ldha, Rest, Pdgfra, Slc16a1, Hk1, and Hk2*) and no change in dedifferentiation markers but restricts β-cell mass expansion. These studies demonstrate that the loss of Tomosyn-2 accelerates the timeline for β-cells to achieve functional maturation and identity, at the expense of β-cell mass expansion.

These results demonstrate that Tomosyn-2 serves as a temporal gatekeeper that coordinates the transition from proliferation to functional maturation in postnatal β-cells. By limiting insulin secretion during early postnatal life, Tomosyn-2 ensures adequate β-cell proliferation to establish a critical β-cell mass required for adult glucose homeostasis. These findings underscore the importance of tightly regulating insulin secretion during the neonatal period to balance β-cell expansion and the acquisition of mature β-cell function.

### Experimental Methods Animals

The blastocysts containing ES cells *Stxbp5l (Tomosyn-2)^tm1a(EUCOMM)Wtsi^* on the JM8A1.N3 were injected into recipient female mice to generate chimeras, which were then crossed with F1 C57BL/6J mice. The presence of floxed alleles was verified using the following primers (FP: TCCTTATCAGCCCACAGCATTGGC and RP: TGAACTGATGGCGAGCTCAGACC), and these mice were backcrossed for 9 generations with the C57BL/6J mice. Tomosyn-2 Flox mice were used to generate *Tomosyn-2^-/-^* mice for this study. Weaning of pups was performed between 3 and 4 weeks of age. All mice had free access to water and a standard chow diet and were housed in a temperature-controlled room with a 12-hour light-dark cycle (6:00 AM – 6:00 PM). All experimental procedures were conducted per the guidelines of the University of Alabama Institutional Animal Care and Use Committee (IACUC-21632).

### Metabolic Phenotyping

In vivo experiments consisted of measuring plasma insulin, blood glucose, and body weight (BW) in *Tomosyn-2^-/-^* and *Tomosyn-2^+/+^*littermate mice. The mice were administered human insulin (Humulin R, Lilly, USA, Cat #0028215-01) at a dose of 0.5 U human insulin/kg body weight (BW) of mice. The dose was administered intraperitoneally following a 6h fast, and the mice were subjected to insulin tolerance testing (ITT). Following a 6-hour fast, the mice were subjected to oral glucose tolerance testing and plasma insulin levels in response to glucose gavage (2 g glucose/kg BW of mice). Blood glucose levels were monitored at various time points by extracting blood from the tail vein and using a glucometer (Contour Next blood glucose meter, Ascensia Diabetes Care, Switzerland) for readings. The plasma insulin content was determined using a mouse insulin ELISA kit from Crystal Chem (USA, Cat #90080).

### Expression constructs

The V5-tagged b-Tomosyn-2 mammalian expression plasmid was subcloned into a Moloney murine leukemia virus-based lentiviral vector (3565) (a gift from Dr. Bill Sugden, University of Wisconsin-Madison, WI)^48^.

### Cell culture and transient transfection

INS1(832/13) pancreatic β-cells (generously provided by Dr. Christopher Newgard, Duke University, NC) were maintained in RPMI 1640 medium supplemented with 10% heat-inactivated fetal bovine serum, 2 mM L-glutamine, 1 mM sodium pyruvate, 10 mM HEPES, 100 U/mL penicillin-streptomycin, and 50 μM β-mercaptoethanol. Cells were seeded at a density of approximately 10,000,000 cells per well in a 10-cm dish. Once the cells reached ∼75–80% confluence (typically the following day), they were transfected with 10 μg of V5-Tomosyn-2-RVV or GFP-RVV (control) using Lipofectamine 2000 (Invitrogen) according to the manufacturer’s instructions. After 36 h, cells were collected for Western blotting or co-immunoprecipitation.

Mouse islets were isolated from 2- to 12-week-old C57BL6/J male or female mice using a collagenase digestion method^59,76^. Isolated islets were cultured overnight in RPMI 1640 medium supplemented with 8 mM glucose. Islets were processed for gene expression analysis and for Western blotting. Further, after 16 h, size-matched islets were hand-picked for a static insulin secretion assay.

### Insulin secretion

Insulin secretion assays using isolated mouse islets were conducted as previously described^48,51,76^. Briefly, six size-matched islets from *Tomosyn-2^-/-^* and *Tomosyn-2^+/+^* mice were manually selected and placed into individual wells of a 96-well plate. The secretion assay was carried out using Krebs–Ringer bicarbonate (KRB) buffer composed of 118.41 mM NaCl, 4.69 mM KCl, 1.18 mM MgSO₄, 1.18 mM KH₂PO₄, 25 mM NaHCO₃, 5 mM HEPES, 2.52 mM CaCl₂, adjusted to pH 7.4, and supplemented with 0.5% BSA. Islets were first preincubated for 45 minutes in 100 μl of KRB buffer containing 2.8 mM glucose. Following preincubation, the buffer was replaced with fresh KRB containing the appropriate insulin secretagogues, and islets were incubated for an additional 45 minutes. The supernatant was then collected to analyze the secreted insulin, and the islets were subsequently lysed in acid ethanol to assess the total insulin content.

Dynamic assessment of insulin secretion was performed using a high-capacity system from BioRep®, which was utilized for islet perifusion as described^16^. A sandwich averaging 75 islets was created in a chamber comprising two layers of Bio-Gel P-2 (Bio-Rad, Cat #1504118) bead solution (200 mg beads/ml in KRB buffer). The chamber temperature was regulated and maintained at 37°C. Islet perfusion was performed at a 200 μl/min flow rate, and secreted insulin was collected in a 96-well plate using an automatic fraction collector. The in-house ELISA quantified cellular and secreted insulin contents. As β-cells are heterogeneous at the postnatal stages, we used all the islets isolated from mice (Supplementary Fig. 1F) for the insulin perfusion data presented in Figure 1 to capture the full potential of insulin secretory capacity.

### Isolation and quantitation of RNA

Total mRNA was extracted from mouse islets by using Qiagen RNeasy Plus Kit (Cat #74034). Following extraction, 1 μg of RNA was used for cDNA synthesis (Applied Biosystems, Cat #74034). The relative mRNA abundance was determined by quantitative PCR using Fast Start SYBR Green (Applied Biosystems) and gene expression was calculated by the comparative Δ*C_t_* method.

### Generating and analyzing bulk RNA-seq data

The total RNAs were extracted from the cell lines as in our prior work^77,78^. The polyadenylated total RNAs were converted into complementary (c)DNA (reads), then processed according to Illumina NextSeq500 guidelines (https://support.illumina.com/ <u>downloads/</u>nextseq-500-user-guide-15046563.html). Then, the raw bulk RNA-seq data were generated and stored in a FASTQ file format. The reads in FASTQ files were trimmed via trim-galore software ^79^; then, via the Burrows-Wheeler Aligner toolki^80^, the trimmed reads were mapped to the Human GRCh38 Reference Genom^81^ to identify the gene corresponding to each read. The percentage of reads that could uniquely map to a single gene in the Human Reference Genome was above 65% (Supplemental Table 1), indicating high data quality. Next, for each gene, the number of reads corresponding to the gene was counted using the HTSeq/0.6.1 package^82^, which provided the initial (non-normalized) gene expression in each replicate. Then, the initial gene expression was normalized, and the Deseq2 toolkit completed the statistical comparison between the *Tomosyn-2^+/+^* and *Tomosyn-2^-/-^*islets^83^. The statistical p-values, computed using DESeq2, were adjusted using the Benjamini^84^ method to address the false-discovery rate issue. Genes with expression magnitude (Deseq2 Basemean) of at least 100, a minimum fold-change of 2, and an adjusted p-value of less than 0.05 were selected as differentially expressed genes.

### Gene ontology, pathway enrichment, and network analysis

Differentially expressed genes from bulk RNA-seq analysis were queried in the DAVID functional annotation tool^85^ to determine which Gene Ontology (GO) terms^86^ and KEGG pathways^87^ were enriched in each cell group. Enriched GO terms and pathways were selected based on two criteria: a) the number of differentially expressed genes in the term/pathway is less than 100; and b) the term/pathway’s Benjamini-adjusted p-value is less than 0.05. Differentially expressed genes belonging to insulin secretion^88^, cell-cycle^89^, and response to endoplasmic reticulum stress (ER-stress)^90^ were queried in the STRING v.12^62^ database to obtain interacting networks among these genes. These networks, including insulin secretion-specific, cell-cycle-specific, ER-stress-specific, and crosstalks among these categories, were visualized via Cytoscape^91^. To determine whether the insulin-secretion genes were more likely to interact with ER-stress or cell-cycle genes, the ratio between the number of insulin-secretion interactions with the other two gene lists and the ratio between the size of the other two gene lists were compared and statistically analyzed via Fisher’s Exact Test.

### Immunoblotting

For protein extraction, mouse islets were isolated from 2-, 4-, 6-, and 15-week-old *Tomosyn-2^-/-^* and *Tomosyn-2^+/+^*mice. Four hundred (per biological replicate) isolated islets were subjected to 400 μl of lysis buffer (1 mM Na_3_VO_4_, 150 mM NaCl, 1 mM EGTA, 20 mM Tris–HCl pH 7.5, protease inhibitor cocktail, 2.5 mM sodium pyrophosphate, 1 mM Na_2_EDTA, 1% Triton X-100, 1 mM β-glyceraldehyde, 1 mM NaF, and 1 mM PMSF). Samples were sonicated for 10 s, and spun at a high speed of 13,000 rpm for 20 min at 4 °C. The supernatant was collected, and the protein concentration was determined using the BCA assay. Protein samples prepared by adding 10 mM DTT were subjected to SDS-PAGE electrophoresis using an 10% polyacrylamide gel^92,93^. The information on the antibodies used for Western blotting is provided in the Supplementary Table 1.

### Embedding human islets for staining

Human islets were bought from the IIDP (Integrated Islet Distribution Program) and processed immediately upon arrival. Islets were handpicked in 1X KRB and then transferred into Eppendorf tubes, where they were washed with 1X PBS. HisoGel™ (LabStorage Systems, Lnc, CAT#HG-4000) aliquots were warmed in a 75°C water bath, and Affi-Gel Blue Media Bead (Bio-Rad, Cat #1537301) aliquots were brought to room temperature. The HisoGel™ and Affi-Gel Blue Media Beads were combined with the islets using a 200 μl pipette and layered on a cold microscope slide in a disc shape to solidify. The disc underwent a series of ethanol and citrisolv washes and was embedded in a paraffin block for sectioning at a thickness of 6 μm, staining, and imaging by confocal microscopy.

### Immunofluorescence

Pancreata from 4- and 6-week-old *Tomosyn-2^-/-^* and *Tomosyn-2^+/+^* mice were fixed in 4% paraformaldehyde at 4°C overnight. The following day, the pancreas tissue was washed with 1X PBS every 10 min for a total of 3 times, subjected to an ethanol gradient, and embedded in paraffin blocks. Tissue sections were cut at a thickness of 6 μm for the entire paraffin block. Slides were then deparaffinized using citrisolv for 5 minutes for a total of 3 times. Rehydration was performed using an ethanol gradient, followed by a 30-minute wash with 1x PBS. Antigen retrieval was performed by placing slides into 1X TEG buffer and subjecting them to power 10 heating in a microwave for 1 minute, followed by power 1 heating for 15 minutes. Once retrieval was completed, the slides were allowed to rest at 27°C for one hour in 1X PBS. Slides were placed into a humidity chamber and blocked with a mixture of 2.5% normal goat serum and 2.5% normal donkey serum for 2 h (2.5% normal donkey serum, 2.5% normal goat serum, 1% BSA, in 1X PBS). Next, the slides were incubated overnight at 4 °C in a primary antibody cocktail. The following day, the slides were washed with 1X PBS as described above, and corresponding secondary antibodies were applied to the slides. They were then incubated in a humidity chamber in the dark for 2 h. Following incubation, the slides were washed again with 1X PBS as described and then mounted with DAPI mounting medium (Southern Biotech, Cat #0100-20). The slides were subjected to confocal imaging using a Zeiss LSM 750 microscope at 20X for mouse islets and 40X magnifications for human islets. The primary antibodies used were anti-Tomosyn-2, anti-Ki67, anti-somatostatin, anti-glucagon, anti-UCN3, and anti-insulin. The secondary antibodies were AF488 anti-guinea pig, AF 647 anti-rabbit, and AF555 anti-mouse. Quantification was performed using Image J. Measures were taken to ensure equal parameters for quantification purposes.

### Proximity Ligation Assay

Proximity ligation assay (PLA) was conducted using the Duolink® In Situ Red Mouse/Rabbit kit (Millipore/Sigma, DUO92101) following the manufacturer’s guidelines. Cells were cultured on poly–D–lysine–coated coverslips and then fixed with 4% paraformaldehyde in phosphate-buffered saline (PBS). Following fixation, samples were blocked in 10% normal donkey serum for 1 hour. After blocking, coverslips were rinsed with PBS containing 0.1% bovine serum albumin (BSA) and 0.01% sodium azide. Cells were then incubated with primary antibodies against Stx1A (mouse monoclonal) and Tomosyn-2 (rabbit polyclonal, Synaptic Systems). The PLA reaction was subsequently carried out per kit instructions. For β-cell identification, a preconjugated Alexa Fluor 488® anti-insulin antibody (rabbit monoclonal, Invitrogen, 53-9769-82) was applied post-PLA. Confocal imaging was performed using a Zeiss LSM710 microscope, and image analysis was conducted with Zen software (Zeiss) for quantifying PLA interaction signals (red puncta) over total nuclei from three independent experiments.

### Islet morphology

Three full-length pancreatic sections (6 μm thick), spaced at least 50 μm apart, were analyzed per sample. Entire sections were imaged at 4× magnification using an Olympus IX81 microscope with stitched image acquisition, followed by analysis in Olympus Cell Sens Dimensions software. Immunostaining was performed using the ABC method with the Alkaline Phosphatase Standard kit (Vector Labs, Cat #AK-5000) in combination with an anti-insulin antibody. Insulin-positive islets were visualized by the presence of blue chromogenic staining (Vector Labs, Cat #SK-5300). Islet morphology was assessed using the manual HSV thresholding feature in Cell Sens software to quantify the following parameters: β-cell cross-sectional area (calculated by multiplying the total β cell cross-sectional area within 3 pancreatic sections/total pancreas area of those 3 sections), circularity (calculated as 4πa/p², where *a* is area and *p* is perimeter), β-cell count per islet, and average β-cell size (determined by dividing total β-cell area by the number of β-cells per islet). β-cell mass was estimated by multiplying the total β-cell cross-sectional area by the weight of the pancreas.

### Proliferation assay

Pancreatic sections were co-stained for Ki67 and insulin to assess β-cell proliferation. Immunofluorescence was performed as previously described. Sections were incubated overnight at 4°C with a primary antibody cocktail containing guinea pig anti-insulin (1:50) and mouse anti-Ki67 (1:500). The following day, slides were washed three times with 1× PBS (10 min each) and then incubated for 2 h at 27°C with secondary antibodies diluted 1:1000: Alexa Fluor 555 anti-mouse and Alexa Fluor 488 anti-guinea pig. Fluorescence imaging was performed at 20× magnification on four pancreatic sections per mouse, each spaced at least 50 μm apart. Ki67⁺β-cells were identified by DAPI-stained nuclei along with staining for insulin and Ki67 immunofluorescent signals. The percentage of proliferating β-cells was calculated as the number of Ki67⁺/Insulin⁺cells divided by the total number of Insulin⁺cells, multiplied by 100.

### Immunoprecipitation

INS1(832/13) cells were plated in 100 mm culture dishes at approximately 60% confluence. For overexpression studies, cells were transiently transfected with 20 μg of either V5-RVV-Tomosyn-2-Ala-RVV or GFP-V5-RVV plasmid using Lipofectamine 2000 (Invitrogen) in Opti-MEM medium, maintaining a 1:1 DNA-to-reagent ratio. After 48 hours, cells were lysed in 1 mL of immunoprecipitation (IP) lysis buffer (25 mM Tris-HCl, pH 7.5; 50 mM NaCl; 2 mM MgCl₂; 1 mM Na₂EDTA; 1 mM EGTA; 1 mM CaCl₂; 0.5% Triton X-100; 2.5 mM sodium pyrophosphate; 1 mM β-glycerophosphate; 1 mM sodium orthovanadate; 1 mM PMSF; and protease inhibitor cocktail). Lysates were incubated on a rotating platform at 4°C for 1 h and then centrifuged at 13,000 rpm for 10 min to separate soluble from the insoluble fraction. The soluble fraction was incubated overnight at 4°C with 2 μg/ml of either anti-V5 or anti-Stx1A antibody, or with an IgG isotype control. The following day, 30 μl of pre-equilibrated magnetic beads were added and incubated for 1.5 hours at 4°C with rotation. Beads were washed three times with 1 ml of IP lysis buffer, and bound proteins were eluted in 40 μl of 2.5× Laemmli sample buffer containing 1 mM DTT. Eluates were heated at 95°C for 5 minutes, separated on 10% or 12% SDS-PAGE gels, and transferred to PVDF membranes. Protein detection was performed using the Clarity Western ECL substrate (Bio-Rad, Cat# 1705061).

### Statistical analysis

Data are represented as means ± SEM. Statistical significance was performed using Student’s two-tailed unpaired t-test for independent data. The significance limit was set at p < 0 .05.

Data availability statement: The data that supports the findings of this study are available in the methods and/or supplementary material of this article. The raw data supporting the conclusions of this article will be made available by the authors without undue reservation.

## Author contributions

Md Mostafizur Rahman, Haifa A. Alsharif, Katherine C. Perez, Justin Alexander, and Sushant Bhatnagar for writing and editing the manuscript; Katherine C. Perez, Justin Alexander, Md Mostafizur Rahman, Haifa A. Alsharif, Yanping Liu, Jeong-A Kim, Chad S. Hunter, Thanh Nguyen, and Sushant Bhatnagar for data curation; Katherine C. Perez, Justin Alexander, Md Mostafizur Rahman, Haifa A. Alsharif for data analysis and data interpretation; Sushant Bhatnagar and Jeong-A Kim provided resources for the project; Sushant Bhatnagar for the conceptualization of the project; Sushant Bhatnagar for the supervision of the project; Sushant Bhatnagar for the funding for the project; Sushant Bhatnagar for project administration. All authors provided critical feedback and contributed to shaping this research, analysis, and manuscript.

## Acknowledgments

We especially thank the human islet donors for their contribution to this study.

