## Supplementary materials and figures for "Tomosyn-2 Regulates Postnatal β-Cell Expansion and Insulin Secretion to Maintain Glucose Homeostasis"

Supplementary figure 1:

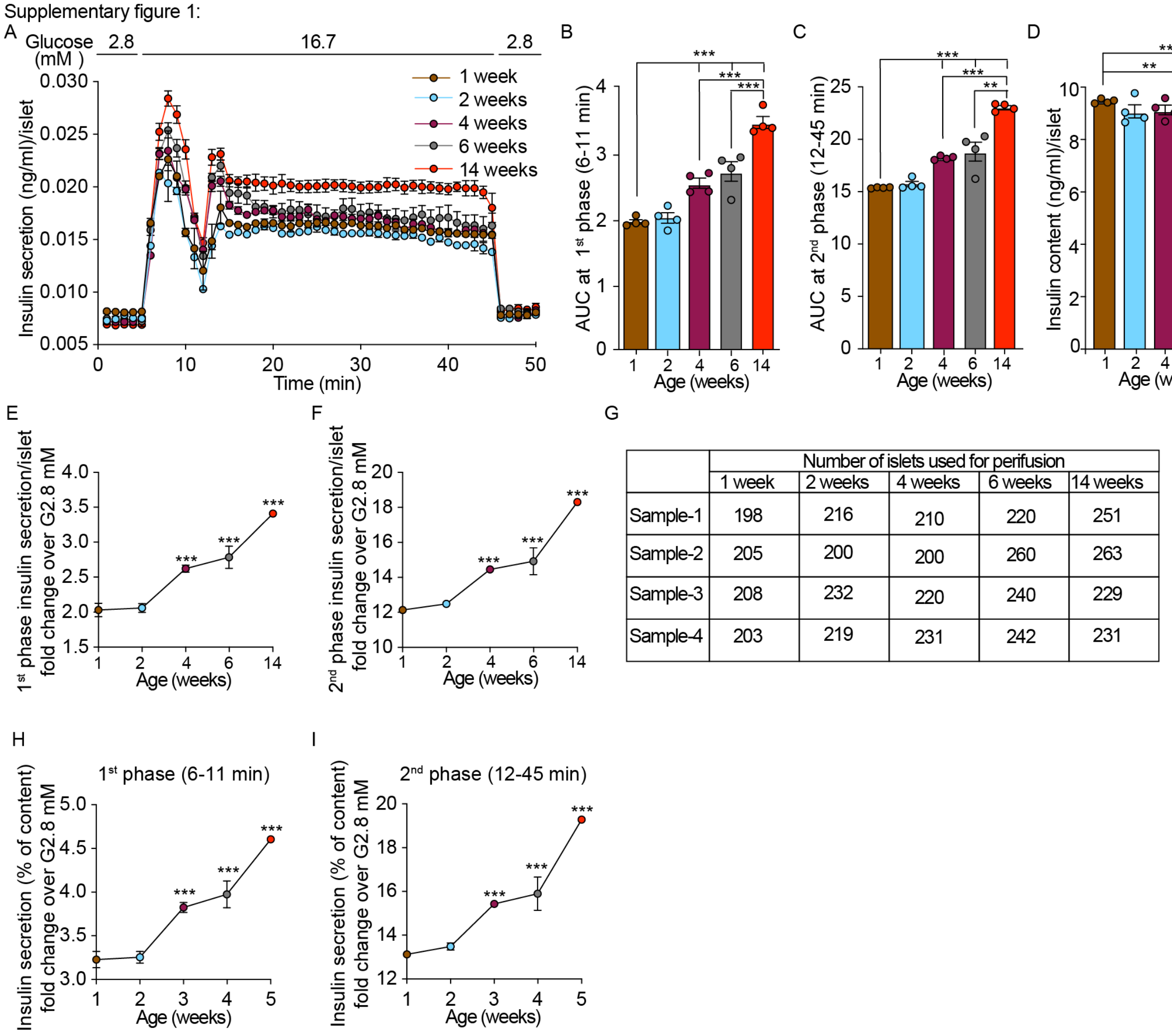

| | Number of islets used for perifusion | | | | |
|---|---|---|---|---|---|
| | 1 week | 2 weeks | 4 weeks | 6 weeks | 14 weeks |
| Sample-1 | 198 | 216 | 210 | 220 | 251 |
| Sample-2 | 205 | 200 | 200 | 260 | 263 |
| Sample-3 | 208 | 232 | 220 | 240 | 229 |
| Sample-4 | 203 | 219 | 231 | 242 | 231 |

**Supplementary Figure 1: Dynamic insulin secretion in islets from postnatal development to adulthood in mice.** (A) Dynamic insulin secretion normalized per islet. (B) Area under the curve (AUC) of first-phase insulin secretion (6–11 min) in response to 16.7 mM glucose (n = 4). (C) AUC of second-phase insulin secretion (12–45 min) in response to 16.7 mM glucose, normalized to basal glucose (2.8 mM) (n = 4). (D) Total insulin content expressed per islet. (E) Amount of insulin secreted during the first phase (n = 4) in response to 16.7 mM glucose. (F) Amount of insulin secreted during the second phase (n = 4) in response to 16.7 mM glucose. (G) The table shows the number of islets used per perifusion chamber in perifusion experiments. All islets from a single mouse were included for insulin secretion measurement, regardless of size matching. Insulin secretion secreted in 1st phase (6-11 min) (H) and 2nd phase (12-45 min) (I) normalized to basal (at 2.8 mM glucose) insulin secretion. Data are presented as mean ± SEMs. ∗P < 0.05, ∗∗P < 0.01, ∗∗∗P < 0.001.

Supplementary Figure 2: ER and mitochondrial stress

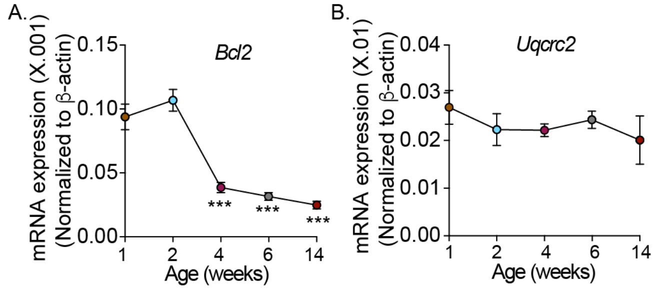

**Supplementary Figure 2. Evaluation of ER stress and mitochondrial gene expression in islets during postnatal development in mice.** (A) mRNA expression of the pro-survival gene *Bcl-2*, normalized to *Actin*, in islets isolated from B6 mice at 1, 2, 4, 6, and 14 weeks of age. (B) mRNA expression of mitochondrial gene *Uqcrc2*, normalized to *Actin*, in islets from B6 mice at 1, 2, 4, 6, and 14 weeks of age. Data are presented as mean ± SEMs.

Supplementary Figure 3

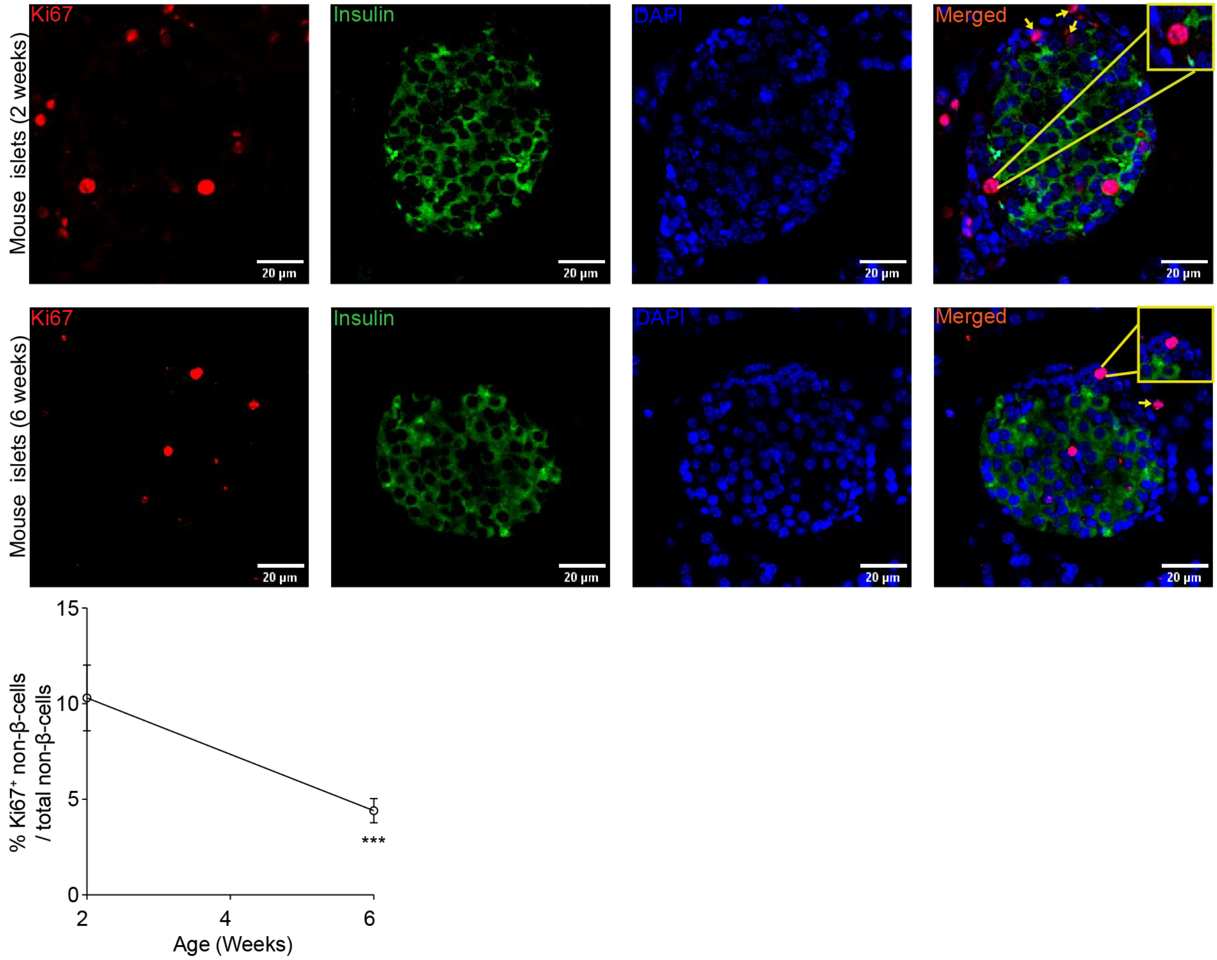

**Supplementary Figure 3. Determination of proliferative non-β-cells in islets during postnatal development in mice.** (A) Immunostaining of islets from B6 mice at 2 weeks (top panel) and 6 weeks (bottom panel) of age showing proliferative cells ($Ki67^+$, red), β-cells ($Insulin^+$, green), and proliferative non-β-cells ($Ki67^+Insulin^-$, indicated with arrows). Images were captured using confocal microscopy at 60× magnification. (B) Quantification of the percentage of proliferative non-β-cells relative to total non-β-cells in islets from 2-week-old and 6-week-old mice (*n* = 4). Data are presented as mean ± SEMs. $*P < 0.05$, $**P < 0.01$, $***P < 0.001$.

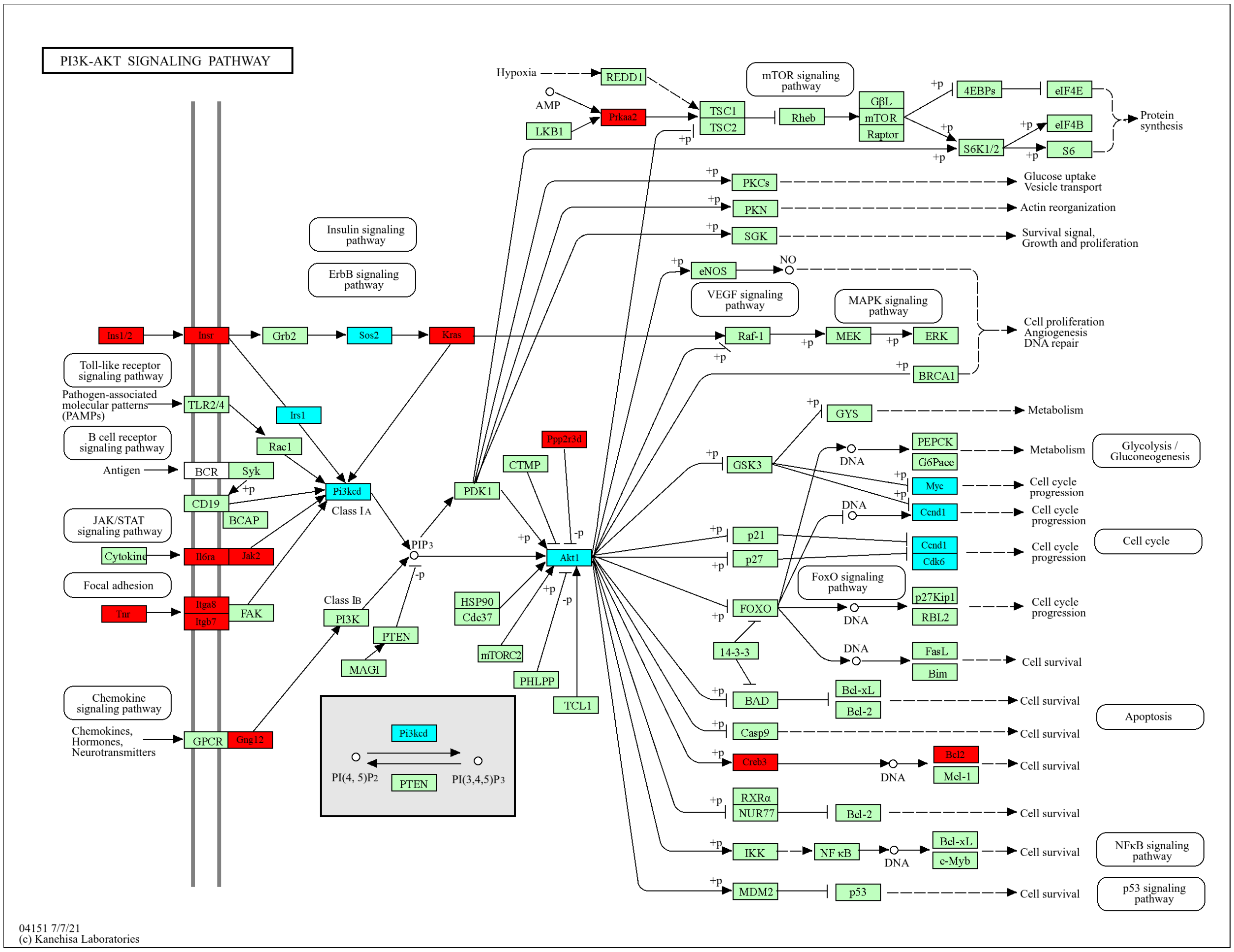

**Supplementary Figure 4. Pi3k-Akt1 signaling pathway.** The diagram of the PI3K-Akt1 signaling pathway was obtained from KEGG (https://www.kegg.jp/pathway/mmu04151). Each gene in the pathway is colored based on the DeSeq2 analysis for differentially expressed genes in *Tomosyn-2*$^{-/-}$ compared to *Tomosyn-2*$^{+/+}$ mouse islets. The upregulated genes are shown in red, downregulated in cyan, and those that are neither upregulated nor downregulated are in gree

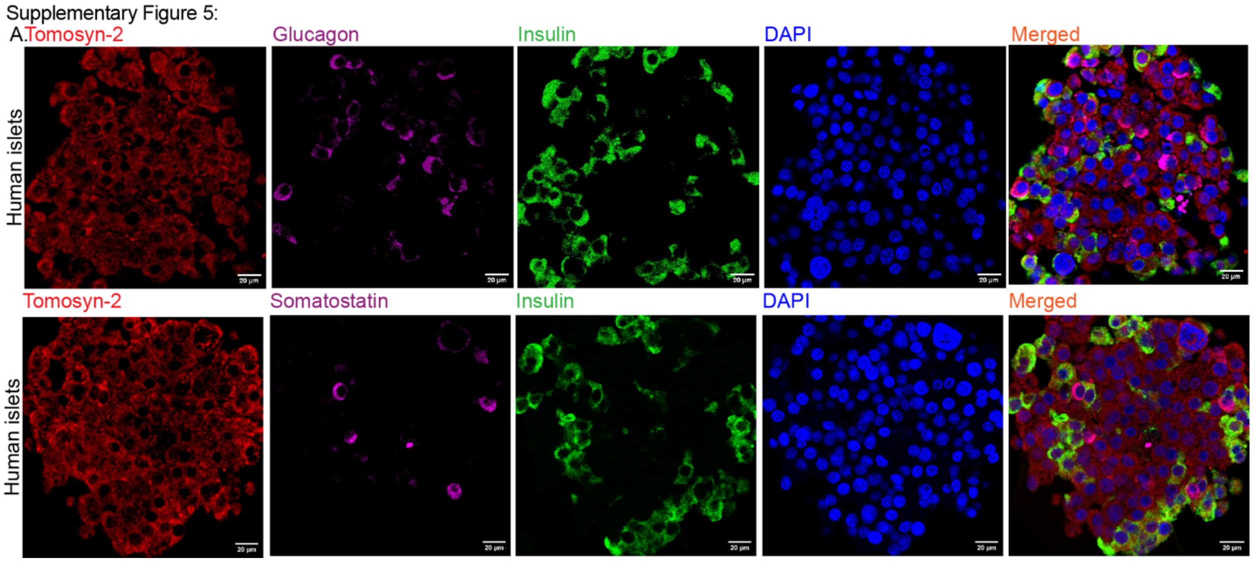

**Supplementary Figure 5. Expression of Tomosyn-2 in non-β-cells of human islets.** (A, top panel) Immunostaining of human islets showing Tomosyn-2 (red), glucagon (purple), insulin (green), and nuclei stained with DAPI (blue). (B, bottom panel) Immunostaining of human islets showing Tomosyn-2 (red), somatostatin (purple), insulin (green), and nuclei stained with DAPI (blue). Images were acquired using a confocal microscope at 60× magnification.

Supplementary Figure 6:

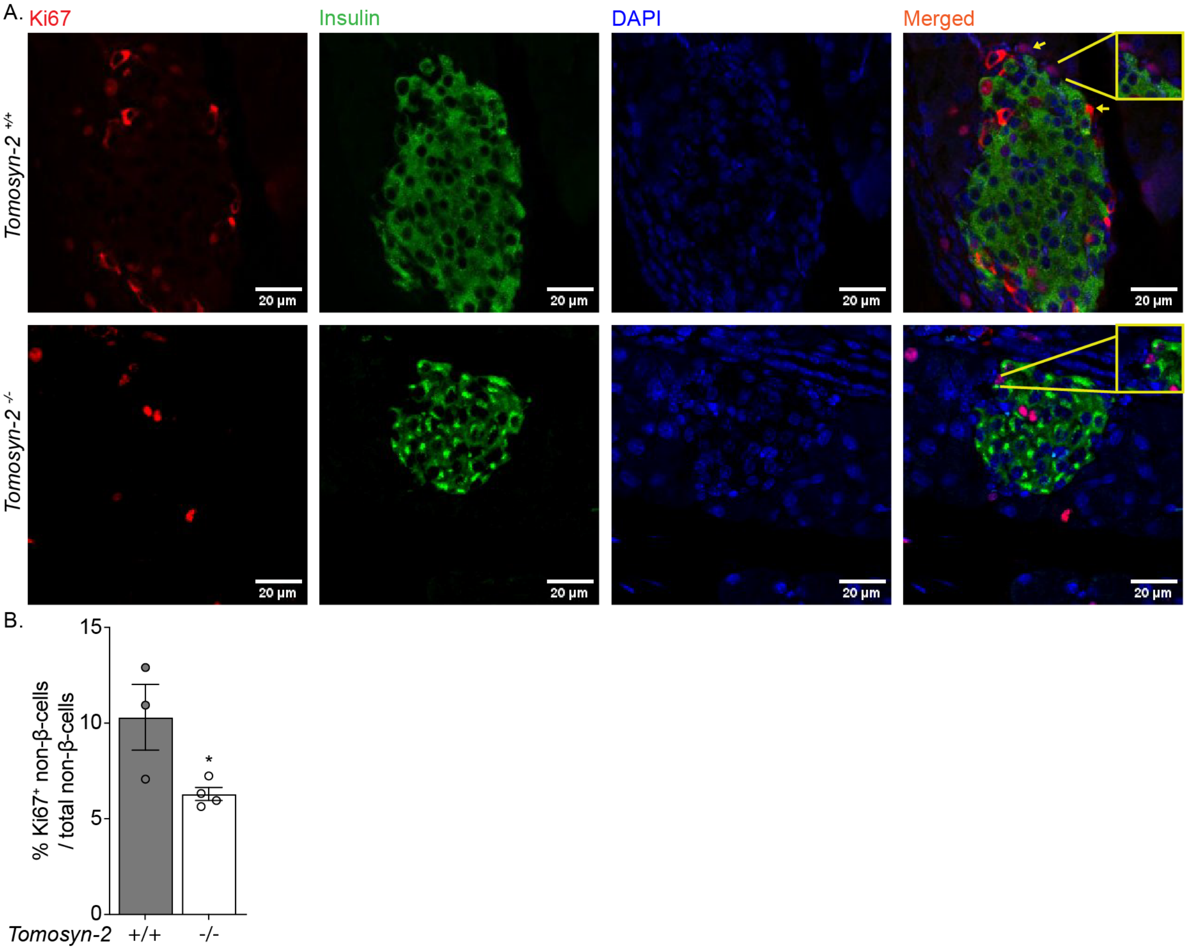

**Supplementary Figure 6. Analysis of proliferative non-β-cells in islets of *Tomosyn-2*$^{-/-}$ and *Tomosyn-2*$^{+/+}$ mice.** (A) Immunostaining of islets showing proliferative cells (Ki67$^{+}$, red), β-cells (insulin$^{+}$, green), and proliferative non-β-cells (Ki67$^{+}$ insulin$^{-}$), indicated with arrows, in islets of *Tomosyn-2*$^{-/-}$ and *Tomosyn-2*$^{+/+}$ mice. Images were acquired using a confocal microscope at 60× magnification. (B) Quantification of proliferative non-β-cells expressed as the percentage of total non-β-cells in islets of *Tomosyn-2*$^{-/-}$ and *Tomosyn-2*$^{+/+}$ mice (n = 3).

**Supplementary Table 1: The list of key reagents used in this study**

| Reagents: | Source: | Catalog No. |
|---|---|---|
| Bio-Gel P-2 | Bio-Rad | 1504118 |
| Applied Biosystems™ High-Capacity cDNA Reverse Transcription Kit | Thermo Scientific | 43-688-13 |
| Collagenase | Sigma | C7657-5G |
| Ficoll® 400 | Sigma | F9378-500G |
| HistoGel | Epredia | HG-4000-012 |
| Humulin R Regular Insulin U | Lilly, USA | 002 8215-01 |
| Insulin Antibody (D3E7 (5B6/6)) - BSA Free | Novusbio | NB100-64697 |
| Rat Insulin Standard | Sigma | 8013-K |
| RNeasy Mini Kit | Qiagen | 74106 |
| Streptaavidine HRP | Thermo Scientific | 21126 |
| Ultra-Sensitive Mouse Insulin ELISA Kit | Crystal Chem, USA | 90080 |
| VECTASTAIN® ABC-AP Kit, Alkaline Phosphatase (Standard) | VectorLabs | AK-5000 |
| VectaMount® AQ Aqueous Mounting Medium | VectorLabs | H-5501-60 |
| Vector® Blue Substrate Kit, Alkaline Phosphatase (AP) | VectorLabs | SK-5300 |

**Supplementary Table 2: The list of antibodies used in this study**

| Antibody: | Source | Catalog No. | Dilution: |
|---|---|---|---|
| Donkey Alexa Fluor 488 | Jackson Immuno Research | 706-545-148 | 1:1000 |
| Glucagon anti-mouse | Sigma-Aldrich | K79bB10 | 1:500 |
| Insulin anti-Guinea Pig | Agilent | IR00261-2 | 1:50 |
| Insulin + proinsulin antibody | Fitzgerald | 10R-I136A | |
| Ki67 anti-mouse | BD Pharmigen | 550609 | 1:500 |
| Mouse Alexa Fluor 647 | Jackson Immuno Research | 715-605-150 | 1:1000 |
| Mouse Alexa Fluor 555 | Jackson Immuno Research | 715-565-151 | 1:1000 |
| Mouse-anti-b- actin | DSHB | 224-236-1 | 1:10,000 |
| Mouse-anti-Syntaxin 1 | Sigma | S0664 | 1:10,000 |
| Normal Donkey Serum | Jackson Immuno Research | 017-000-121 | 2.5% |
| Normal Goat Serum | Jackson Immuno Research | 005-000-121 | 2.5% |
| Peroxidase Goat Anti-Rabbit IgG (H+L) | Jackson Immune Research | 111-035-003 | 1:10,000 |
| Rabbit Alexa Fluor 647 | Jackson Immuno Research | 111-605-003 | 1:1000 |
| Rabbit- anti-Vamp2 | Synaptic System | 104 008 | 1:3000 |
| Somatostatin anti-mouse | Santa Cruz Technology | SC74556 | 1:500 |
| Tomosyn-2 anti-rabbit | Cedarlane Labs | 183203(SY) | 1:100 |

**Supplementary Table 2: qPCR primers**

| qPCR Primers | Forward (5'-3') | Reverse (5'-3') |
|---|---|---|
| Aurkb | CTTCTACGACCAGCAGAGGATC | GGCATCTGACAGTTCCTCCATG |
| β-actin | TGTGATGGTGGGAATGGGTCAGAA | TGTGTTGCCAGATCTTCTCCATGT |
| Bcl-2 | ATGCCTTTGTGGAACTATATGGC | GGTATGCACCCAGAGTGATGC |
| Ccnd1 | TCTACACCGACAACTCCATCCG | TCTGGCATTTTGGAGAGGAAGTG |
| Foxm1 | CACTTGGATTGAGGACCACTT | GTCGTTTCTGCTGTGATTCC |
| Glut2 | CAGTTCGGCTATGACATCGGT | GTTAATGGCAGCTTTCCGGTC |
| Glp1R | TGAACCTGTTTGCATCCTTCA | ACTTGGCAAGCCTGCATTTGA |
| Ins1 | CCAGCTATAATCAGAGACCA | CCAGGTGGGGACCACAAAGA |
| Ins2 | GGCTTCTTCTACACACCCAT | CCAAGGTCTGAAGGTCACCT |
| Kcnj11 | TGTGCAGAATATCGTCGGGCTGAT | GCATGCTTGCTGAAGATGAGGGTT |
| Ki67 | TCCTTTGGTGGGCACCTAAGACCTG | TGATGGTTGAGGTCGTTCCTTGATG |
| Nkx6.1 | CCTCTGGACCCGAACTCTGA | GCTGCCACCGCTCGATT |
| Tomosyn-2 KO | CACTGCATTCTAGTTGTGGTTTG | GCAAACCGGGAATCTGGATAA |
| Tomosyn-2 W | CCAGCTTGTTACTGTCACTATAGG | CATGCCGAACTGTGTGAAAGAGAG |

Supplementary Table 3

### Checklist of human islets used in this research.

Adapted from Hart NJ, Powers AC (2018) Progress, challenges, and suggestions for using human islets to understand islet biology and human diabetes. Diabetologia https://doi.org/10.1007/s00125-018-4772-2

| Islet preparation | 1 | 2 | 3 | 4 |
|---|---|---|---|---|
| **MANDATORY INFORMATION** | | | | |
| Unique identifier | RRID:SAMN44571547 | RRID:SAMN38428316 | RRID:SAMN39671770 | |
| Donor age (years) | 49 | 51 | 19 years | |
| Donor sex (M/F) | Male | Male | Male | |
| Donor BMI (kg/m$^2$) | 24.70 | 25.60 | 31.70 | |
| Donor $HbA_{1c}$ or other measure of blood glucose control | 5.0 | 5.9 | 5.0 | |
| Origin/source of islets[b] | IIDP | IIDP | IIDP | |
| Islet isolation centre | Prodo lab | Wisconsin | Pennsylvania | |
| Donor history of diabetes? Please select yes/no from drop down list | No | No | No | |
| **If Yes, complete the next two lines if this information is available** | | | | |
| Diabetes duration (years) | | | | |
| Glucose-lowering therapy at time of death[c] | | | | |

*Continues on the next page*

| RECOMMENDED INFORMATION | | | |
|---|---|---|---|
| Donor cause of death | Cerebrovascular/stroke | Anoxia | Head trauma |
| Warm ischaemia time (h) | *No* | Yes | No |
| Cold ischaemia time (h) | *18 hours 20 minutes* | 4 hours 45 minutes | 18 hours 20 minutes |
| Estimated purity (%) | 95% | 95% | 85% |
| Estimated viability (%) | 95% | 98% | 89% |
| Total culture time (h)[d] | 1 days 23 hours. | 2 days 18 hours. | 1 days 23 hours |
| Glucose-stimulated insulin secretion or other functional measurement | Glucose Stimulated Insulin Release Stimulation Index (SI): 7.7 | Data not available | 5.6 |
| Handpicked to purity? Please select yes/no from drop down list | | | |
| Additional notes | | | |

[a]If you have used more than eight islet preparations, please complete additional forms as necessary
[b]For example, IIDP, ECIT, Alberta IsletCore
[c]Please specify the therapy/therapies
[d]Time of islet culture at the isolation centre, during shipment and at the receiving laboratory
